# A signature of deep credit assignment in the human Orbitofrontal cortex

**DOI:** 10.64898/2026.11.20.767874

**Authors:** Shany Grossman, Loek Van Rossem, Noa L. Hedrich, Andrew Saxe, Nicolas W. Schuck

## Abstract

Humans are remarkably efficient at learning from limited experience because they can generalize quickly. Doing so, however, requires first learning about the latent structures that govern outcomes. The dominant theoretical framework to understand this interactive cycle between learning task representations and generalisation is deep learning, which updates parameters throughout multi-layered neural networks using deep credit assignment. Yet, whether the brain uses a similar “deep” approach to learning is unclear. Here, we derive and test a prediction that uniquely characterizes deep credit assignment, namely, that a single learning experience should trigger immediate, structured representation changes that are non-local (i.e. generalize beyond the observed experience), and adapt as learning proceeds. Using fMRI (N=51) and neural network simulations, we tested whether feedback on a limited set of items induces structured changes in the representations of other, non-feedbacked items during a partially-observable reinforcement learning task. We find that with sufficient learning, feedback on a small set of trials begins to induce rapid, context-dependent shifts in non-local orbitofrontal cortex representations that extend beyond experienced items and predict concurrent improvements in behavioral generalization. The neural findings and their link to behaviour were mirrored by representation learning dynamics in performance-matched recurrent neural networks, trained using gradient descent on the same curricula as participants. These findings reveal fast, non-local dynamics of representation learning that support generalization and are shared across humans and distributed deep networks.

## Introduction

Faced with ever-changing streams of rich sensory inputs, humans must balance their capacity to remember individual events with the ability to generalize past experiences to novel situations^1,2^. Generalization is thought to be rooted in learning task representations that emphasize task-relevant features and de-emphasize idiosyncratic features with no predictive value^3–5^. Humans, for instance, will naturally learn that restaurants in one neighborhood tend to be expensive while those in another are affordable, which allows them to infer affordability of never-visited restaurants, depending on their location but independently of other aspects. Yet it remains poorly understood how the process of learning task-appropriate representations, i.e. the emphasis on neighborhoods in our example, interacts with the process of generalization from one observation to another.

The ability of modern deep learning systems to learn task representations that support generalization is considered central to their success^6–8^ and is thought to make neural networks a powerful model for brain function^9–12^. Like the brain, neural networks rely on distributed processing, which allows them to capture latent structure through the degree of similarity between representations^13,14^. While comparisons between networks and brains have mostly focused on the structured representations that result from learning, little attention has been paid to the learning process that produces them, and how it interacts with generalisation in deep neural networks. In artificial neural networks, learning proceeds via deep credit assignment, which uses deviations between the output and the correct answer to update all weights across multiple layers of representation, typically via gradient descent. A core implication of deep credit assignment over distributed representations is that individual representations are difficult to manipulate independently: weight updates following even a single training example immediately and inevitably alter hidden representations and outputs for many other inputs. The precise structure of such rapid alterations, which we call non-local updates, is critical for learning success, as it determines whether changes lead to overgeneralization or catastrophic forgetting^2^, or whether they enable effective generalization of knowledge. This makes reconciling local and non-local learning a fundamental challenge for both brains^15–17^ and deep artificial neural networks^18,19^. By analytically deriving learning dynamics in linear networks trained with gradient descent, we show that the structure of non-local updates to hidden representations can adapt to learned task structure only if credit assignment is *deep*, i.e. operates over at least two hidden layers (see Methods, ‘Analytic derivation of non-local updates in deep linear networks’). Non-local updates that (a) occur immediately with each experience and (b) adapt over time are thus a signature of deep credit assignment. Our main question was therefore whether this signature of deep credit assignment can be found in the brain, and where.

In the brain, converging evidence suggests that the orbitofrontal cortex (OFC) has a pivotal role in processing reward, as well as in representing latent, inferred structures in humans^20–24^, non-human primates^25,26^ and rodents^27–32^ (see^33,34^ for recent reviews). Other work has suggested an essential role of the hippocampus in representing latent structures^35,23,36,37^. Most of these studies measured existing representations of previously learned task rules, rather than probing their structured dynamics as they are learned in real-time, with some notable exceptions^38–40^. Hence, we hypothesized that OFC, and possibly the hippocampus, are shaped by deep credit assignment, and should therefore show the signature described above.

Here, we use analytical derivations, simulations and neuroimaging to investigate such rapid representation learning dynamics in the context of human reinforcement learning. Specifically, we ask how reward feedback on a small set of trials immediately generalizes to representations of, and responses to, distinct future observations. Using multivariate analyses of neuroimaging (fMRI) data, we find that abstract task representations (i.e., task states) emerge slowly in the OFC over the course of learning, accompanied by structured non-local updates dominated by unobservable context identity, as predicted by a deep credit assignment account. These context-dependent non-local updates were predictive of subsequent performance improvements on non-feedbacked items, linking neural and behavioral generalization. Neural and behavioral findings were mirrored by representation learning dynamics found in performance-matched recurrent neural networks, trained using gradient descent on the same curriculum as participants. In contrast, neither visual regions nor the hippocampus showed the predicted pattern or the same relation to behavior. Taken together, these findings demonstrate a previously unexplored neural mechanism for facilitating representation learning in the human brain.

## Results

Data from 51 participants (mean age 26 ± 4.5, 36 females) who completed a 2-session value-based reinforcement learning task while being in the fMRI were included in all analyses (out of 62 scanned, 10 did not reach performance criteria and one excluded due to excessive motion, see Methods). During the experiment, participants took on the role of a realtor who had to learn the house valuations of a set of clients from feedback. The task included eight different clients and eight different houses (i.e., 64 client-house valuations). However, a latent outcome structure meant that instead of learning 64 distinct values, participants could learn only 8 different values and generalize these across different client-house pairs that differed only in irrelevant features (participants knew about the existence of the structure, but had to learn which client/house features were relevant and how). Concretely, the house values depended on only some, but not all features of the client and only some, but not all features of the houses: although clients differed in whether they wore glasses or not, whether they were man or woman, and their specific facial features, only one binary feature (eye-sight / gender) was relevant for their house preference. Similarly, while houses had three binary features (location, material, exterior facility), only two of these features determined the outcome with one of them being more important than the other (relevant client and house features were counterbalanced across participants). The 8 sets of client-house combinations that share relevant features (but can differ in irrelevant features) are henceforth labeled as task states (see Fig. 1b, labeled as a1-a4 and b1-b4).

**Figure 1.**
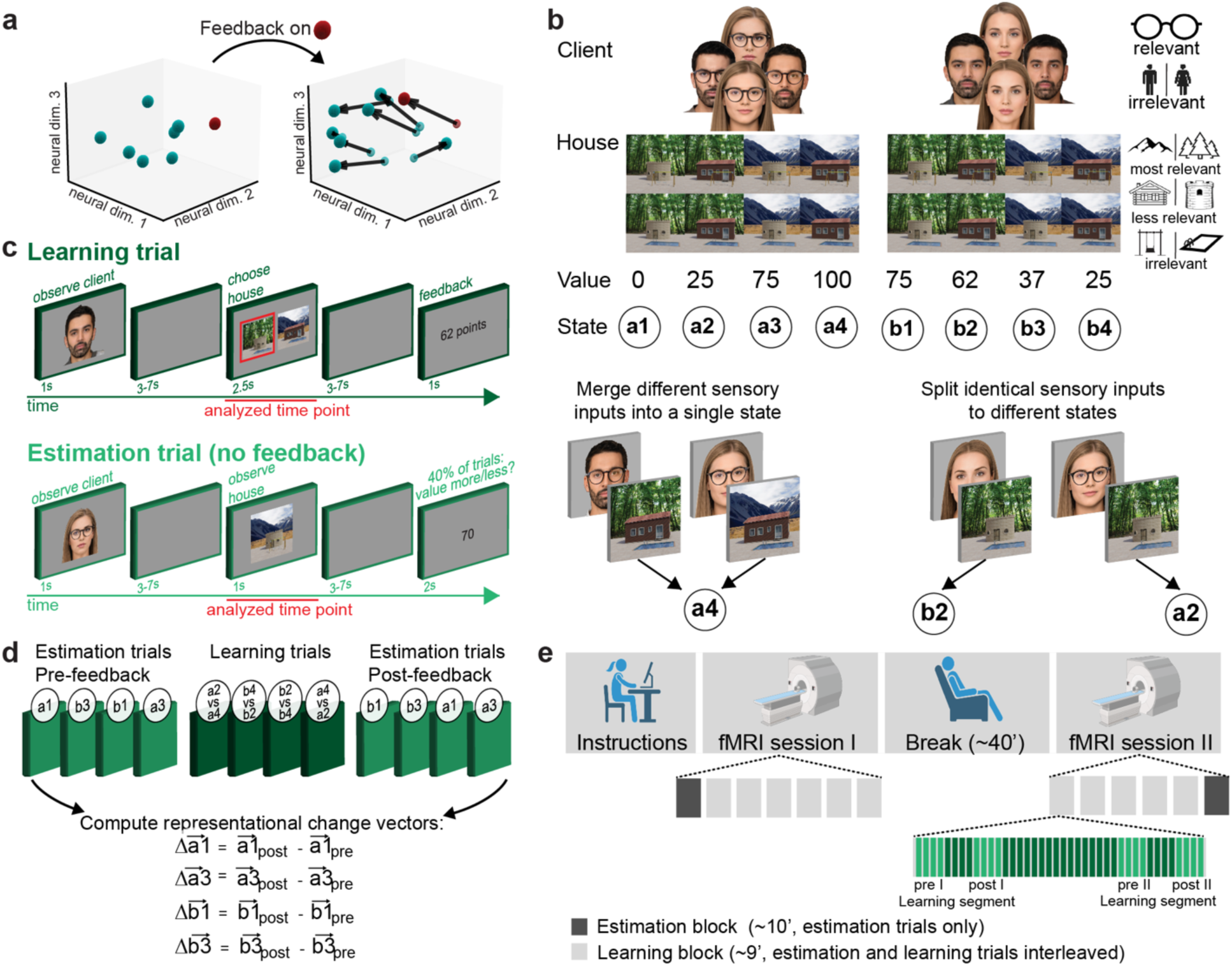
Research question and experimental design. **a.** Illustration of non-local representation learning. Feedback on a single decision concurrently shifts representations of other task items, depicted as points in neural space. These non-local shifts are structured by learned task knowledge and can emerge only in deep networks (>1 hidden layer), as derived for gradient descent learning (see analytic derivations in Methods). **b.** Example value function used in the task, defined over eight latent states. Here, client type was defined based on whether the client wore glasses or not, and house type was defined as a combination of the house location (mountain / forest) and its material (wooden / stone). Gender, specific facial identity and house exterior (pool / swing) were irrelevant. Collapsing across irrelevant variations compressed the 64 stimuli combinations into 8 meaningful latent states (a1-a4, b1-b4) with their assigned values. A second solution, with relevance assigned to different features, was counterbalanced across participants (see Methods). **c.** Schematic of trial types. In learning trials (top), participants viewed a client followed by two houses and were asked to select which of the houses the client would prefer. Decisions were followed by numerical feedback indicating the value of the chosen house. In estimation trials (bottom), participants viewed a client followed by a single house and were instructed to think about the corresponding value. Forty percent of estimation trials were followed by a value probe that asked whether the current client-house pair is worth more or less than a presented threshold. No feedback was provided. **d.** Schematic of a single learning segment embedded within pre-and post-estimation segments. Estimation trials included only latent states that have not been feedbacked during the interleaving learning segment, enabling us to measure non-local shifts in emerging task representation. **e.** The experiment consisted of two subsequent fMRI sessions. Participants completed 11 learning blocks and two additional estimation blocks.

A second major aspect of the value function was that client identity flipped the house valuations (e.g., clients with and without glasses had opposite preferences), while house features acted additively on the value. The client was therefore the most impactful aspect of the task, making the learning problem non-linear and solvable only by non-linear neural networks with at least one hidden layer.

Performing the task involved learning and estimation trials. On learning trials (Fig. 1c, top), participants first viewed an image of a client (1 second) followed by two simultaneously presented houses (2.5 seconds), and then had to decide which of the two houses to offer the client. Each choice was followed by a reward indicating the value of the chosen house for the client (0-100 points). In estimation trials (Fig. 1c bottom), participants were presented with a client followed by a single house and were asked to think about the value associated with that client-house pair. Participants’ value estimate was probed in 40% of estimation trials, but no feedback was provided (see Methods). The two trial types were intermixed such that short segments of four learning trials (∼1 minute of learning) were preceded and immediately followed by 4-5 estimation trials (hereafter: pre- and post-estimation trials, see Fig. 1d, details see methods), which, crucially, only involved states that were not presented in the interleaving learning trials. Since participants could not learn from estimation trials, and they involved states not observed in the interleaving learning segment, the comparison of pre-versus post-estimation trials allowed us to precisely measure non-local representational change as indexed by changes in BOLD activation patterns of task states from pre to post estimation trials.

A task block consisted of two learning segments, each surrounded by pre- and post-estimation segments, which will be the major focus of our analyses. To ensure sufficient learning, the two pre-post measures around small learning segments were separated by an additional 16 learning trials in the middle of each block (Fig. 1e). Overall, the task consisted of 11 blocks, resulting in 264 learning and 198 estimation trials (24 and 18 learning and estimation trials per block, respectively). In addition, participants completed two blocks consisting of only estimation trials, at the start and end of the experiment. Fig. 1e depicts the experimental procedure in detail.

### Behavioral modelling: participants learn the compressed task states

Choice accuracy increased from 59% in block 1 to 93% in block 11 for learning trials and from 62% to 85% for estimation trials, indicating learning of the client-house value mapping (Fig. 2a, paired t-tests block 11 vs. 1; *t*s(50) = 14.4 and 8.9, respectively, both *ps* < .001). To test whether participants learned to generalize by compressing the 64 client-house pairs into the 8 latent states, we fitted two linear function approximation RL models^41^ to choice data: a perceptual model that assumes independent value learning of 64 client-house pairs, and a compressed model that assumes value learning in a hand-crafted latent space of only the 8 true states (Fig. 2b). Using a standard delta learning rule, both models can learn the values of each input, although the compressed model learns faster: its reduced state space means each observed outcome automatically generalizes to seven other client house-pairs mapped onto the same state.

**Figure 2.**
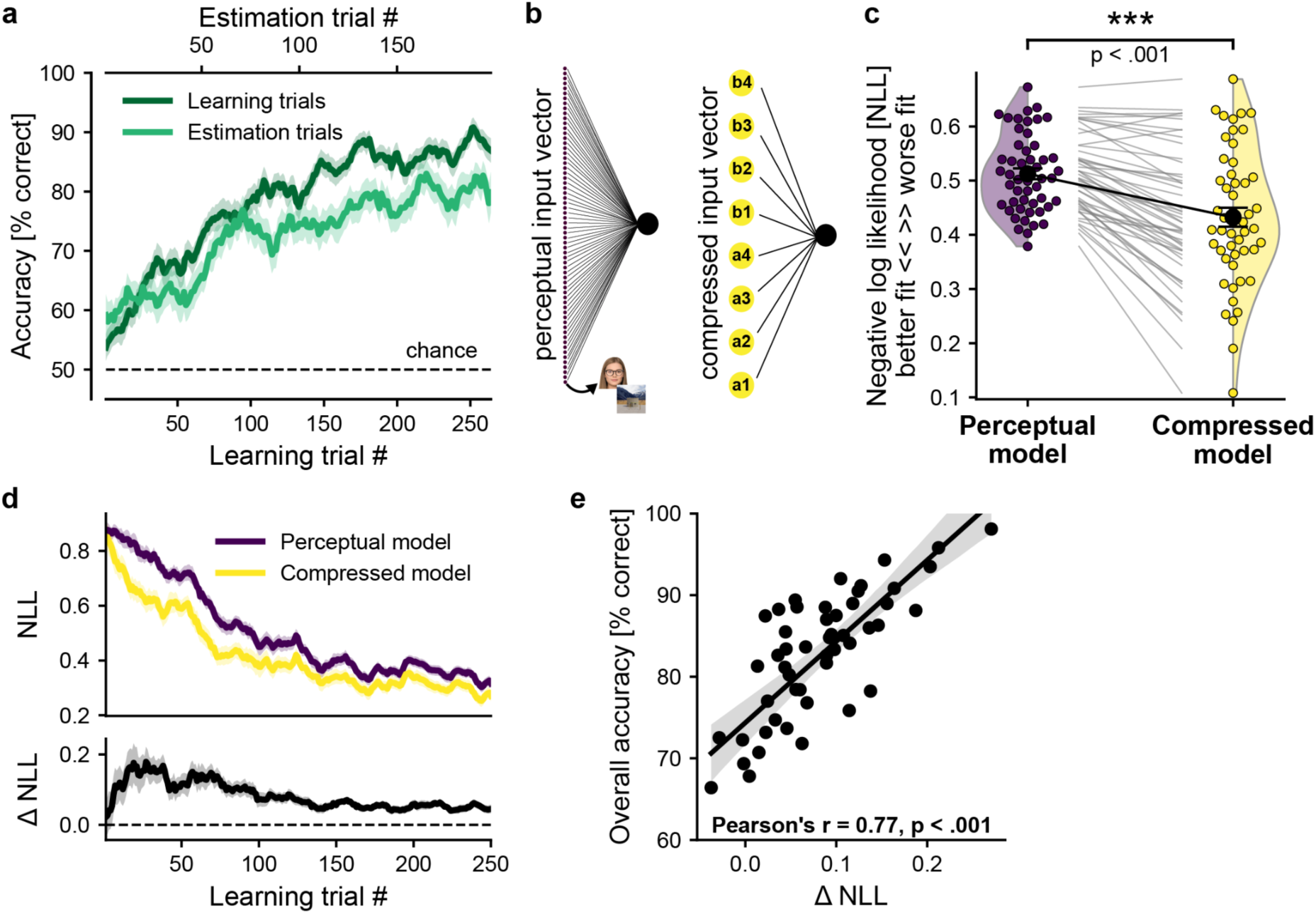
Behavioral modelling demonstrates participants learn the compressed task structure. **a.** Mean accuracy curves of all participants (N=51) in learning and estimation trials. **b.** Schematic of the 2 linear approximation models fitted to choice data. Inputs in the perceptual model were 64 one-hot vectors indexing unique client-house stimuli pairs. In the compressed model, inputs were encoded as 8 one-hot vectors indexing the 8 latent task states. **c.** Model fits for individual participants, quantified as the mean negative log likelihood (NLL) across all learning trials. Each circle denotes a participant, with lines connecting individual participants. p value is derived from a paired t-test. **d.** Top: single-trial NLL over the course of learning for the two models, averaged across participants. Bottom: mean difference between the two time-series presented on top, wherein positive values reflect a better fit to the compressed model. All shaded areas denote SEM. **e.** Scatter plot showing the advantage of the compressed over the perceptual model (x-axis) against overall learning accuracy (y-axis). Greater relative fit of the compressed model is significantly associated with better performance. Black line and shaded area denote linear fit and 95% confidence interval, respectively.

The compressed model yielded a significantly better fit to participants’ choices (paired t-test on mean negative log likelihood: t(50) = -8.9, p < .001, Fig 2c). Observing the trial-by-trial time series of model fits (Fig. 2d) showed that the advantage of the compressed model emerged quickly and was most substantial during the first half of learning, indicating that participants inferred the hidden structure of the task and leveraged it to expedite learning. As expected, the relative advantage of compressed over perceptual model fit was positively correlated with overall accuracy (Fig. 2e; Pearson’s r = .77, p < .001).

### Task state representations emerge in the Orbitofrontal cortex with learning

A core prediction of a deep credit assignment account is that emergence of structured latent state representations leads to fast non-local learning. Having established that learning and generalization behavior was guided by inferred latent task states, we therefore next asked whether latent state representations emerged in neuroimaging data over time. We focused on the Orbitofrontal cortex (OFC) and the Hippocampus, which have both previously been implicated in the representation of latent task states ^21,31,40^. In addition, we investigated two functionally-defined high-order visual regions implicated in processing our visual stimuli, the ventro-temporal visual cortex (VTC) and the Fusiform face area (FFA). Activation patterns for all fMRI analyses were quantified as single trial betas elicited at the time of the house pair / single house presentation in learning / estimation trials, respectively (see Methods). Note that while the client was no longer displayed during the analyzed time window itself, it was part of the hypothesized task states, rendering our task partially-observable.

We used a cross-validated representational similarity (RSA) approach to test which ROIs show evidence for learning the representations of partially-observable latent task states. A linear mixed-effects model asked whether activity patterns became more similar for trial pairs involving the same latent states than for pairs that did not, increasingly over time (first vs. second half of the experiment, henceforth early vs. late). Because identical latent states could map onto different visual inputs (with different irrelevant features), and identical visual inputs could map onto different latent states (when trials differed by the no longer shown relevant client feature) this analysis was unconfounded by sensory input (Fig. 1b, bottom). Outcome similarity was controlled for by adding value similarity as a confound factor. Lastly, task-irrelevant similarity was modelled using a factor which indexed the ratio of identical irrelevant features across the two trials. The core effect of interest was the interaction of task phase with latent state encoding, capturing learning-related increases in encoding relevant but not irrelevant information. The same analysis was applied to learning and estimation trials (for details, see Methods).

The results of this analysis are shown in Fig. 3. In both trial types, the OFC showed significant main effects for state (Learning: β_state_=.008±.001, z_state_=8.7, p<.001; Estimation: β_state_=.006±.001, z_state_=5.3, p<.001) and value (Learning: β_value_=.007±.001, z_value_=6.2, p<.001; Estimation: β_value_=.003±.001, z_value_=2.1, p=.03), but no evidence for encoding irrelevant task features (all ps > .06). We also found an interaction between learning phase and state (Learning: β_state:phase_=0.005±0.001, z_state:phase_=5.7, p<.001; Estimation: β_state:phase_=0.003±0.001, z_state:phase_=3.0, p<.01), reflecting an increase in state encoding from early to late learning, as well as specificity to task-relevant features, even after accounting for value effects. Refining the task phase regressor from a binary early/late to successive block pairs afforded higher temporal resolution and revealed an increasing effect of state, which became significant at block pair 3-4 for learning trials and at block pair 5-6 for estimation trials (Fig. 3a, bottom row; see Fig. S1 for remaining ROIs).

**Figure 3.**
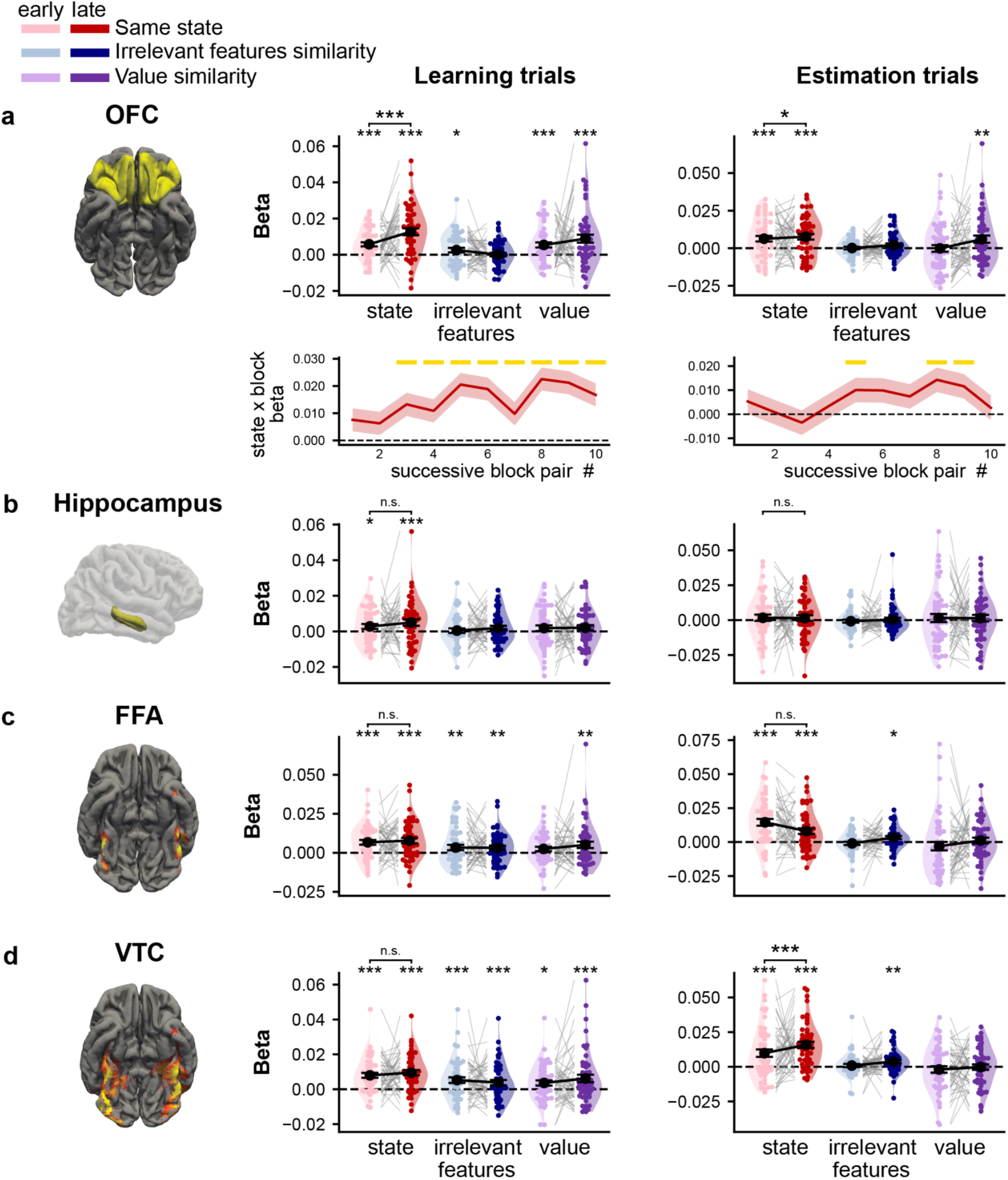
Emergence of latent task states in OFC over the course of learning. **a.** Top: Results from a linear-mixed effects model of cross-run single-trial activation patterns’ correlations in the OFC. Significance of single learning phases (early / late) was derived from modelling each learning phase in isolation. For data visualization purposes, individual dots represent betas derived from single-participant OLS models. Bottom: Time series of same state coefficients relative to the first cross-run data point. The first coefficient reflects the state effect for block 1–2 cross-run correlations, whereas subsequent coefficients indicate the relative increase in state encoding for later block pairs. **b–d.** Same analysis as in a (top), applied to Hippocampus, FFA and VTC ROIs. All error bars and shaded areas indicate standard error of the mean. FFA and VTC ROIs were functionally defined in individual brains, based on an independent visual localizer scan. For visualization purposes, anatomically-masked group contrast maps are presented here, arbitrarily thresholded at t(50) > 4.

The same analysis for the hippocampus, FFA and VTC showed that no other region had a significant interaction of state encoding with learning phase in both learning and estimation trials. All 3 ROIs showed a main effect for state in both trial types (all ps<.001), with the exception of estimation trials in the hippocampus (p=0.5). Unlike OFC, however, VTC and FFA showed significant main effects for encoding irrelevant features similarity (ps < 0.018), suggesting that the state effect of these visual regions is at least in part due to input similarity. A significant interaction of state with learning phase was observed only for estimation but not learning trials in the VTC, and in none of the trial types in the FFA. For brevity and compactness, statistics for all ROIs and trial types are reported in Table S1.

Taken together, these findings demonstrate the emergence of latent state encoding in the OFC as learning unfolds. Evidence from other ROIs suggests state and value signals are widespread in the brain, but only the OFC has both a specificity to relevant signals and a learning related dynamic in both trial types, which will be the core focus of this study.

### OFC exhibits dynamic non-local updates as predicted by deep credit assignment

We next turned to our primary research question: does learning of task states induce adaptive non-local updates of task representations? To answer this, we extracted the activity patterns of all estimation trials before and after a given learning segment, and computed the pre-post representational differences for each unique estimation trial (Fig. 1d). Because the absolute amount of change between single trials is highly susceptible to instantaneous noise sources, we focused on directional changes and asked whether the change vectors were structured in a non-random manner through an RSA approach. Specifically, we tested whether the direction of change (in high-dimensional activity space) in each representation was linked to how other representations changed in a systematic manner. For every segment, we first computed above-mentioned pre-post differences and then calculated a set of 6 pairwise correlations between change vectors, which we term (non-local) co-updates. As previously mentioned, the states for which change vectors were computed in each segment were never presented during the interleaving four learning trials and therefore provided a measure of fast non-local representational shifts in the absence of direct feedback. Based on theoretical insights about the signature of deep credit assignment in neural networks, our primary hypothesis was that OFC would show a temporally dynamic change in the correlation structure of change vectors (i.e. co-updates) that increasingly reflects learned task dimensions.

The primary form of correlation structure we examined was whether context identity shaped non-local representation learning. As before, task-context was defined by the relevant client feature presented before the house display which was the analyzed time window. For each learning segment, we computed the mean co-update (correlations between change vectors) among states sharing vs. not sharing the same context, and averaged these across all 22 learning segments. OFC showed a significant context effect (Fig. 4b), i.e., non-local same-context co-updates were significantly greater than zero (1-sample t-tests, same context: t(50)=2.9, p=.005, Cohen’s d=0.4) and significantly larger than co-updates between states from different contexts (paired t-test: t(50) = 2.8, p = .007, Cohen’s d = 0.4). Different-context co-updates, in contrast, were smaller than zero (t(50) = -2.2, p = .03, Cohen’s d = -0.31). A time-series analysis of context-dependent co-updates showed that this effect only emerged with time in OFC (Fig. 4c). Hence, the characteristics of OFC representational generalisation dynamics during learning matched the conceptual predictions of a deep credit assignment mechanism in deep (>1) layers of neural networks. The Hippocampus, FFA and VTC, showed no evidence for context-dependent co-updates (all *p*s >.1).

**Figure 4.**
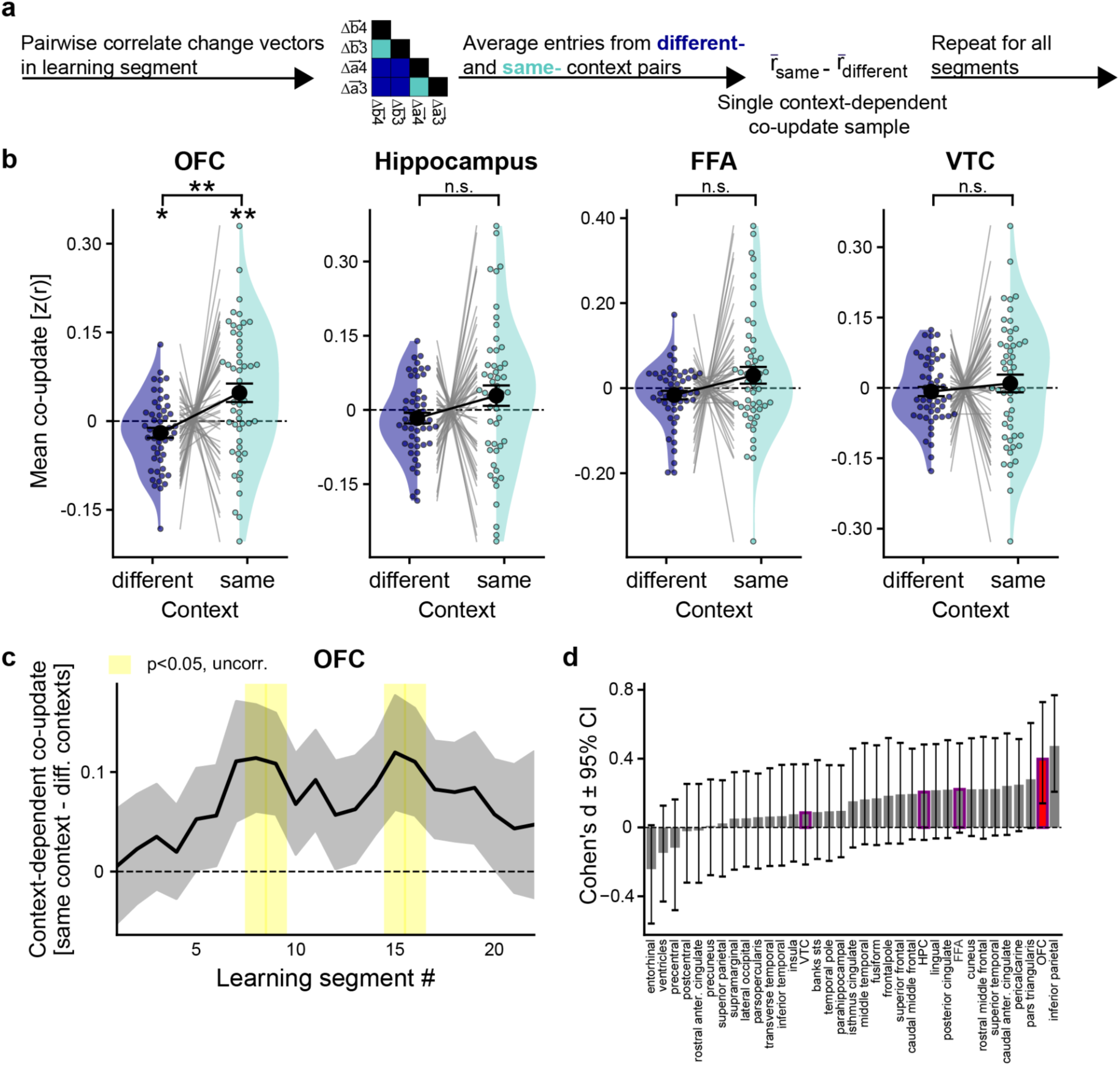
OFC shows context-dependent non-local representation learning. **a.** Schematic of the analysis carried on change vectors to extract a measure of context-dependent co-updates in a single learning segment. All single co-update values were z-scored within each participant and ROI, then averaged within each learning segment for each participant. **b.** Mean co-update values between change vectors originating in the same versus different contexts. OFC showed a significant context-dependent co-update effect. Individual significance levels per class (same / different contexts) were derived from 1-sample t-tests. Difference between conditions derived from a paired t-test. Error bars denote SEM. **c.** Time-series of context-dependent co-updates, averaged across participants and smoothed with a running average of 4 learning segments. Shaded area denotes SEM, yellow patches denote p<.05 (1-sample t-test, uncorrected). **d.** An exploratory analysis depicting the effect-size of mean context-dependent co-update across all anatomical ROIs derived from the Desikan–Killiany atlas freesurfer parcellation. Error bars denote bootstrapped 95% confidence intervals (1000 bootstraps). Bars stroked in purple mark the a-priori ROIs analyzed, OFC additionally colored in red.

To assess the spatial selectivity of the effect, we ran an exploratory analysis of context-dependent co-update effects in all 31 anatomical ROIs in the Desikan–Killiany freesurfer cortical parcellation ^42^, as well as in the ventricles. The results showed that the effect in OFC was among the strongest found across the entire brain (Fig. 4c), with only the inferior parietal lobule showing an effect size greater than the OFC. Inspecting the time-series of context-dependent co-updates in the inferior parietal lobule showed the effect in this region emerges later relative to the OFC, towards the end of learning (Fig. S2). We also confirmed the robustness of the context effect in the OFC by performing robustness tests in which we computed the effect size for different processing choices, including noise modelling, spatial smoothing, dropout threshold, correlation metric and matched samples size of same- and different-context pairs per learning segment (Fig. S3a).

Next, we asked whether the structure of co-updates in OFC also reflected a second core dimension of the learned task structure: value. Given that cross-context comparisons were more widely sampled per learning segment, we focused on cross-context co-updates, making this effect independent of our previous analysis. We compared cross-context state pairs with maximal vs. minimal value differences within a segment and averaged this across all learning segments. We again found an effect in OFC, wherein states from different contexts diverged more strongly when their values were closest than when they were furthest apart (Fig. 5a, one-sample t-test of co-updates for minimal value difference: t(50) = -3.2, p = .002, Cohen’s d=0.28; paired t-test between co-updates for maximal vs. minimal value difference: t(50) = -2, p = .04, Cohen’s d = 0.28). This is consistent with representational separation compensating where value least distinguishes states from different contexts. Lastly, we tested whether the OFC effect was independent of visual similarity by comparing the cross-context co-updates of states sharing the same vs. different house type (i.e., the same two relevant house features, Fig. 5b). OFC co-updates were not significantly influenced by visual input similarities (all ps >.06). Interestingly, however, HPC showed a significant divergence between cross-context states sharing the same visual input but pertaining to different unobservable contexts, echoing and extending upon previous findings about context-dependent repulsion of hippocampal representations for visually similar inputs^43^ (Fig. S4b, one-sample t-test of co-updates for same visual input: t(50) = -3, p = .004, Cohen’s d=-0.4; paired t-test between co-updates for same vs. different visual input: t(50) = 2.5, p = .01, Cohen’s d = 0.35). Taken together, these results provide evidence for dynamic non-local generalization in OFC that was specific to latent task aspects and independent of sensory similarities, in line with mechanisms of deep credit assignment.

**Figure 5.**
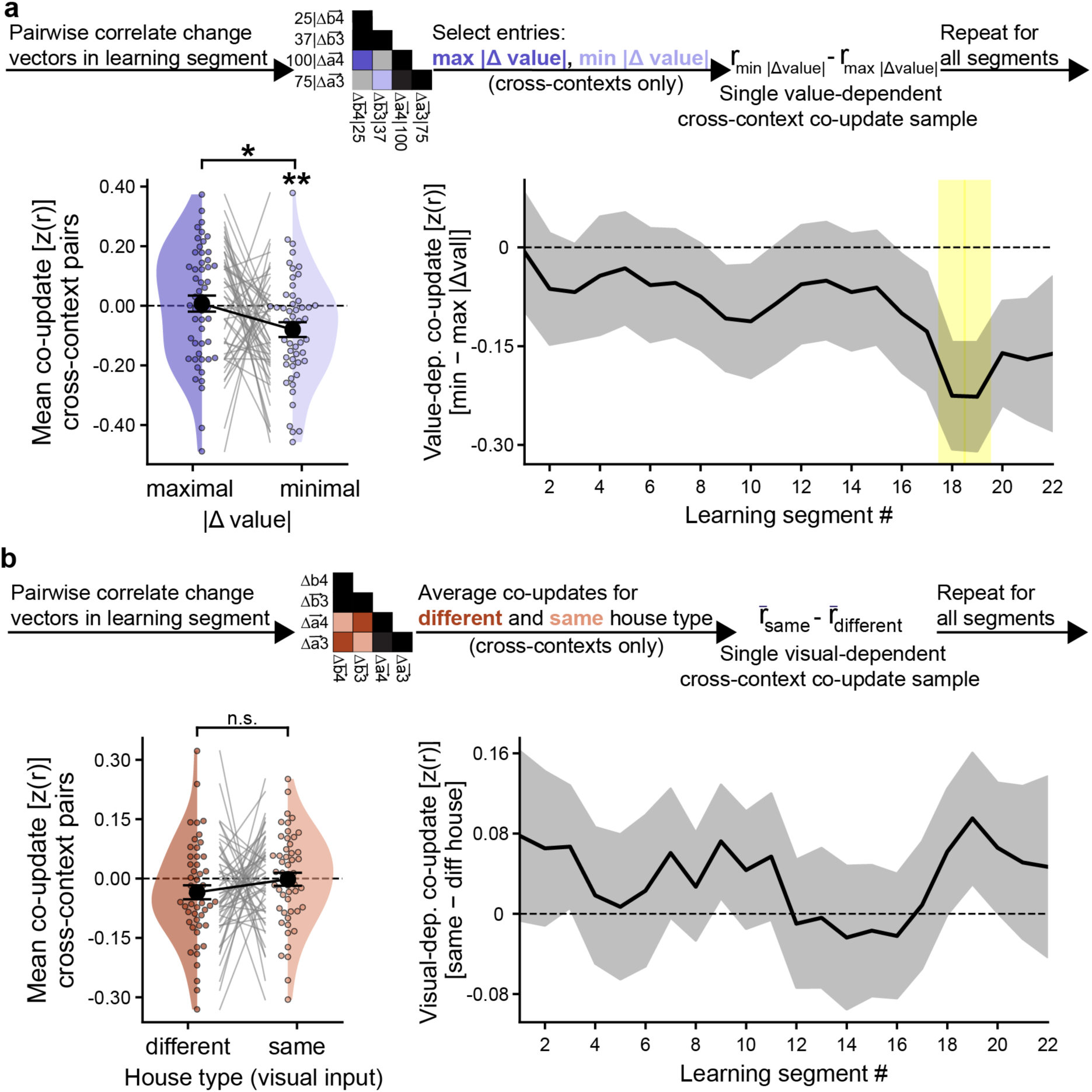
Cross-context non-local representation learning in the OFC reflects learned values, but not visual similarity. **a.** Top: scheme for computing a value-dependent co-update in a single learning segment. Bottom left: mean non-local co-updates for pairs with maximal vs. minimal value difference. Cross-context states with similar values (i.e., minimal value difference) significantly diverge representation space. Bottom left: temporally-resolved time series of value-dependent co-updates across learning. **b.** Top: scheme for computing a visual-dependent co-update in a single learning segment. Bottom left: mean non-local co-updates for pairs with different versus same visual input (i.e., house type). Bottom right: temporally-resolved time series of visual-dependent co-updates across learning. OFC did not show a significant effect. Error bars denote SEM. Significance derived from 1-sample and paired t-tests for single condition and cross-condition comparisons, respectively. *: p<.05, **: p<.01.

To test whether the non-local effects could arise from unsupervised mechanisms, independent of the local feedback received in learning trials, we repeated the analysis on final estimation blocks (see Methods). These blocks followed all learning blocks and contained only estimation trials, with no interleaved learning trials. OFC showed neither a context-dependent co-update (t(50) = 0.36, p = .72, d = 0.05, BF₀₁ = 6.2) nor a value-dependent co-update (t(50) = 0.62, p = .54, d = 0.08, BF₀₁ = 5.5) in these blocks, with Bayes factors providing moderate evidence for the absence of both effects. Thus, the non-local co-updates observed during learning were not present in the absence of interleaved learning trials, suggesting that they are driven by local feedback rather than by unsupervised dynamics, such as consolidation.

### Non-local context-dependent representation learning in the OFC is linked to generalization

We next asked whether the neural signature of deep credit assignment in OFC was related to participants’ generalization. To extract maximally specific behavioral characterisations, we computed the difference in accuracy between probed pre- and post-estimation trials for each segment capturing performance changes in estimation trials that cannot be attributed to feedback and hence present genuine generalisation. In the OFC, estimation probes with a positive generalization index (better performance after than before) were accompanied by significantly larger context-dependent co-updates, as compared with estimation probes with negative generalization index (Fig. 6a, one-sample t-test on mean co-updates for positive generalization: t(50)=3, p=.003, Cohen’s d=0.43; paired t-test positive vs. negative generalization: t(50)=2.7, p=.01, Cohen’s d=0.38). This effect remained significant across all processing variants tested in a robustness analysis (all *ps* < .05; all Cohen’s ds > 0.3; Fig. S3b). Co-updates of estimation probes with saturated performance (100% accuracy) before and after learning did not differ significantly from zero (p=.2). We next separated the context-dependent effect into its two constitutive components, (i) a cross-context push reflecting representations from different contexts diverging in representational space, and (ii) a within-context pull, reflecting correlated change directions of representations sharing the same context. Repeating the analyses described above separately for each neural component showed that cross-context push was significantly negative in segments characterized by behavioral generalization (Fig. 6b, one-sample t-test: t(50) = -2.2, p = .03, Cohen’s d = 0.43) and marginally smaller compared to no generalization segments (paired t-test: t(50) = -1.8, p = .08, Cohen’s d = 0.25). Within-context pull was significantly greater than zero in generalization segments (Fig. 6d, one-sample t-test: t(50) = 2.7, p = .009, Cohen’s d = 0.39) and significantly greater compared to no generalization segments (paired t-test: t(50) = 2.4, p = .018, Cohen’s d = 0.35).

**Figure 6.**
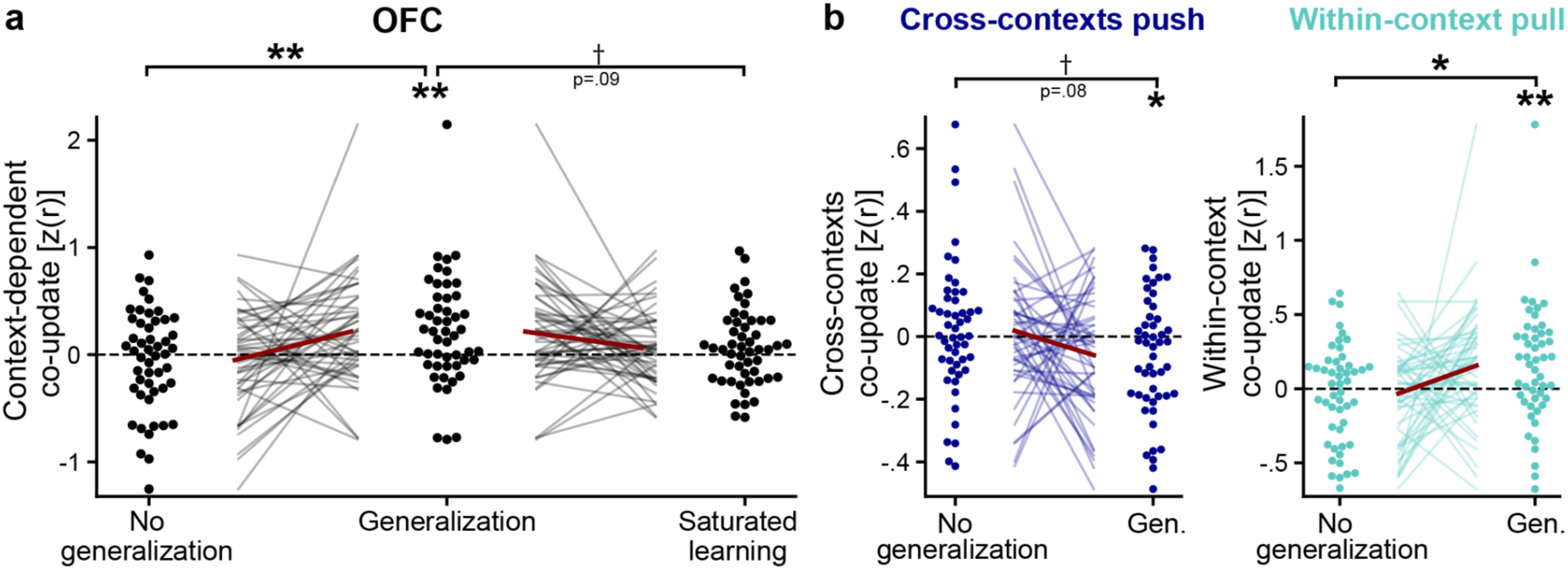
OFC context-dependent non-local co-updates linked to participants’ generalization. **a.** Mean OFC context-dependent co-updates categorized by accuracy change across pre- and post-estimation trials probes: post relative to pre performance increase (generalization), decrease (no generalization), and stable ceiling performance (saturated learning). Significance levels between conditions were derived from paired t-tests, and per condition from 1-sample t-tests. Learning segments with generalization were associated with significantly stronger context-dependent co-updates relative to no generalization segments. **b.** Same as a, applied separately to the two neural components constituting context-dependent non-local updates: (i) Cross-contexts push (divergence between cross-context states) and (ii) within-context pull (common directional shift between same-context states).

A between-subject analysis found that participant-wise average magnitudes of OFC context-dependent co-updates did not significantly correlate with interindividual differences in overall learning performance, indexed by the mean accuracy in all learning trials (Pearson’s r = -.2, p = .15). The link between co-updates and behavior thus appears to hold within rather than between individuals, possibly because our performance-based inclusion criterion limited variability in overall performance. The same analyses of non-local co-update signals in the remaining ROIs (HPC, VTC and FFA) and the Inferior Parietal Lobule (which showed a large context-dependent co-update effect in our whole brain exploratory analysis) did not reveal any significant relation to local generalization indices or to overall accuracy (all ps > .1, see Fig. S2 for the IPL).

### Performance-matched deep neural networks show non-local context-dependent representation learning, similar to OFC

Our research question was motivated by the observation that linear neural networks trained with gradient descent exhibit non-local co-updates with only 2 or more hidden layers (see Methods: Analytic derivation of non-local updates in linear networks). To confirm this prediction in non-linear networks trained on the same curriculum, we set up a recurrent neural network able to perform our task. The model received binary input vectors across two time-steps that reflected the client and house presentations in our task, respectively, and was trained to predict the true outcomes using online gradient descent along a mean-squared error loss function. Inputs were first passed to a simple recurrent layer (RNN, necessary to model our partially-observable 2-step task), followed by a fully connected (FC) hidden layer and an output node (Fig. 7a, see Methods for details). Fifty-one networks were trained on the exact same learning curricula as our 51 participants and fit to participants’ choices through optimizing the learning rate, and the size of initialized weights in each of the hidden layers (see Methods for details), yielding one matched network per participant (see Fig. 7b for average performance, and Fig. S5 for individual curves). Although some differences in learning speed persisted, matched networks reached the same performance level as humans when trained on the same curriculum, as evidenced by statistically indistinguishable accuracy by the end of learning (paired t-test between networks’ and participants’ accuracies in last three learning blocks: t(50)=−1.75, p= .09).

**Figure 7.**
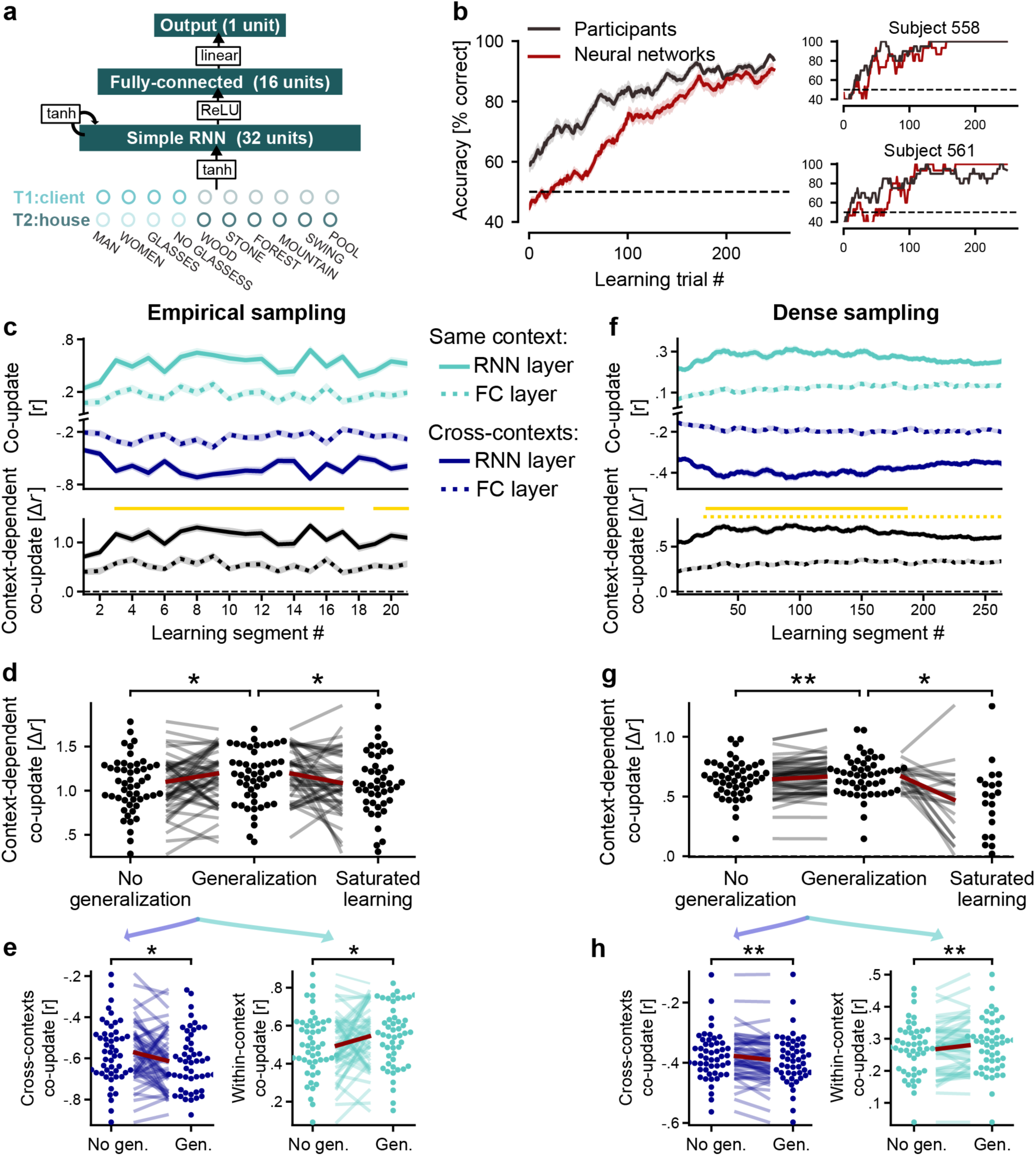
Context-dependent co-updates in performance-matched networks and their link to generalization. **a.** Scheme of the network architecture trained on the same task and curricula as participants. On each learning trial, the network predicted the value of the left and right house. Weights were updated on each trial according to the feedback received on the chosen house. **b**. Mean learning curves of participants and their performance-matched networks (N=51) and 2 individual examples. **c.** Non-local context-dependent co-updates in both hidden layers (RNN, FC) of the networks (N=51). Top: separated time-series for same and cross-context pairs. Bottom: delta time series (same minus different). Yellow lines denote a significant increase in context-dependent co-update relative to the first learning segment (paired t-tests across networks, FDR-corrected at α=.05 within each layer; solid: RNN layer, dotted: FC layer). **d.** Context-dependent co-updates in the RNN layer categorized by performance change in single learning segments, equivalent to Fig. 6a. **e.** The same analysis as in d, applied separately on the two activity components constituent of context-dependent co-update, equivalent to Fig. 6b. **f-h.** The same analysis as in panels c-e, only that context-dependent co-update and generalization indices were sampled densely, for every learning trial and for all items unobserved on that trial.

Applying the non-local representation learning analysis to hidden activations of performance-matched networks revealed, in the RNN layer, context-dependent co-updates that gradually increased over learning and returned to baseline towards its end (Fig. 7c), similar to the time series observed for the OFC (Fig. 4b). Although the effect significantly increased over time, it appeared to be present at the very first learning segment. Repeating the analysis for the irrelevant context feature showed a comparable onset effect followed by a significant decrease (Fig. S6), indicating that the onset effect reflects input overlap, whereas the increase was specific to task features the network had learned to be relevant, in line with theoretical predictions.

We also asked whether the observed pattern could arise with Hebbian learning, which does not rely on deep credit assignment principles. Training neural networks with Oja’s rule ^44^ did not reach human-level performance (mean accuracy in the last learning block: 69.8% ± 12.2% SD). Further, their non-local co-updates emerged early, remained constant, and crucially were equally present for relevant and irrelevant context features (Fig. S7). This is to be expected from a rule that uses no output targets, as the relevant and irrelevant context features are similar in terms of input structure and only appear distinct given their targets.

Context-dependent co-updates in the RNN layer had the qualitatively same relations to behavior as observed in the OFC (Fig. 6), i.e. co-updates were significantly larger in generalization vs. no generalization segments (Fig. 7d, paired t-test: t(50) = 2.2, p = .03, Cohen’s d = 0.3). Moreover, context-dependent co-updates were less pronounced for saturated learning segments (Fig. 7d, paired t-test generalization vs. saturated learning in RNN layer: t(45) = 2, p = .05, Cohen’s d = 0.3). A follow up analysis with increased statistical power, using denser performance and co-update sampling of neural networks, revealed identical effects with increased effect sizes (Fig. 7g; paired t-test for generalization vs. no generalization in RNN layer: t(50) = 3, p = .004, Cohen’s d =.4; paired t-test for generalization vs. saturated learning in RNN layer: t(19) = 2.6, p=.016, Cohen’s d = 0.59). Robustness analysis showed that an intra-subject link to generalization was present in various held out initializations of the networks, but only when using the dense sampling approach described above. Non-local learning in the FC layer was not linked to instantaneous generalization segments for either the empirically-matched sampling or for the dense sampling (all ps >.18).

The neural networks did not capture all aspects of our data. In particular, unlike OFC, non-local co-updates in the networks were partly driven by input similarity, as shown by effects already present at learning onset for pairs that consistently shared a context feature, whether relevant or irrelevant (Fig. 7c,f; Fig. S6). Consistent with this, identical house inputs significantly reduced repulsion between cross-context pairs (paired t-test across networks: t(50)=11.2, p<.001, Cohen’s d=1.6). These results indicate that learning in OFC is better able to override input similarity relative to the simple networks we considered. Because the networks could not fully override input similarity, and cross-context pairs with maximal value differences typically share the same house in our design, our networks were poorly suited for testing purely value-dependent non-local co-updates (as carried out on BOLD signals in Fig. 5a).

## Discussion

Learning latent structures of the environment that are not observable from immediate sensory inputs is fundamental to making goal-directed decisions in complex and ever-changing environments. A key benefit of having such structural knowledge is that it can guide future learning by determining how feedback is generalized. This generalization will not only update how agents value and respond to related items, but also restructure the underlying internal representations across multiple levels. In other words, latent knowledge is not only reflected in representational structure, but also in how feedback adjusts the entire representational organization far beyond the observed event.

Here, we provide an analytical derivation of deep credit assignment in linear networks trained with gradient descent, and show that such adaptive representational generalization based on learned latent structure can occur only in deep architectures with at least 2 hidden layers, thus constituting a hallmark of deep credit assignment. Using neuroimaging, we provide evidence that the human OFC not only represents learned latent task structures, but also dynamically generalizes local feedback to update non-locally learned latent task representations, consistent with a deep credit assignment account. Specifically, representational generalization occurred after only 4 learning trials and primarily reflected a learned context feature that was absent from the screen throughout the analyzed time period, with representations from the same context shifting in tandem, and those from different contexts shifting in opposing directions. Fluctuations in this co-updating across learning segments were significantly linked to a behavioral generalization immediately afterwards. Neural networks trained on the same task and curriculum as participants confirmed our analytical predictions and showed the same link to generalization. Hippocampal and visual brain regions, however, did not show compatible patterns.

OFC co-updates were not explained by (fixed) visual similarity or unsupervised learning rules. First, co-updates only emerged over the course of learning and were attenuated when performance was saturated. Second, they were not present in the final blocks without feedback, which speaks against unsupervised dynamics, such as consolidation. Lastly, control analyses showed that cross-context co-updates were shaped by learned values, but not by a-priori visual similarity.

Our findings align with and extend previous findings that implicated the OFC in representing values^45,46^ as well as a cognitive map of the task^27,47,35^. They also build on evidence that prediction errors drive trial-by-trial changes in OFC activity patterns that relate to behavioral adaptation ^39^, and show that such feedback-driven updates extend non-locally to representations of other items, structured by latent task dimensions. Our findings are also in line with the general idea that the brain might employ a form of deep learning^9,11,48^, and the specific suggestion of OFC reflecting late hidden representations of a deep RL architecture^10,33^.

The fact that performance-matched networks showed the same non-local updates and link to generalization as OFC fits with theoretical work on how deep networks learn. Prior theoretical work has shown that deep learning in the ‘rich’ regime involves learning structured task-specific representations that impart inductive biases which shape generalization^49,50^, and which share universal aspects across many network architectures^51^. Our current findings suggest this process is underpinned by non-local updates, present in both deep layers of neural networks and in the human OFC. Neural networks did not capture all aspects of our data, however. Non-local representational shifts in our performance-matched networks remained sensitive to input similarity, whether task-relevant or not, while OFC showed no advantage for cross-context pairs sharing the same house inputs over those that did not. Learning in OFC therefore seems better able to override input similarity than the simple networks we considered, possibly suggesting that the networks we used where not deep enough. Future work should investigate which settings promote an OFC-like invariance to input overlap and whether such invariance accelerates learning in artificial systems.

An essential question raised by our findings is how the brain implements deep credit assignment, since exact backpropagation is considered biologically implausible ^52^. We therefore deliberately tested a signature that does not require exact gradients or backpropagation, only that error signals change synapses at more than one processing stage. This is also the case in more plausible approximations such as feedback alignment or predictive coding^53^. In essence, our findings therefore suggest that the brain assigns credit to “deep” abstract representations in OFC, independently of how this assignment is achieved.

Another open issue is how the here described findings interact with other mechanisms that could have similar effects, such as feedback-driven attentional changes^54^, or replay. Feedback can for instance trigger replay of task states that were not observed^55–57^, and reward can spread to never-rewarded items through hippocampal reactivation of associated memories^58^. While the absence of non-local co-updates in blocks without feedback argues against unsupervised mechanisms, such feedback-triggered processes could still play a role in our task. Future studies could therefore test whether non-local updates follow single feedback events, and whether they co-occur with replay.

A third set of open questions concerns the roles of other brain regions. An exploratory whole brain analysis showed that the inferior parietal lobule exhibited context-dependent co-updates with an even larger effect size than OFC. In this region, however, the effect emerged later and was not related to behavioral generalization. Given the role of posterior parietal cortex in abstract task representations^59–61^, this might be worth investigating further. The hippocampus did not show context-dependent non-local co-updates, but visually identical items from different contexts were pushed apart. This is in line with previous findings of a repulsion effect in human hippocampus, wherein representations of visually similar inputs are more strongly decoupled when linked to distinct broader contexts^62,43^, and with findings showing that hippocampal task representations retain specific sensory instantiations of latent structures^31^, while those in OFC/vmPFC generalize over them^63^. Visual regions also encoded task states, but at least partly because of input similarity, since both VTC and FFA also encoded irrelevant features, and no learning-dependent increase was found. It is possible that our task has involved visual reinstantiation of previously observed relevant visual information (i.e. the relevant client feature). With regards to value signals, recent studies in both humans and rodents have found evidence for value signals in somatosensory regions which are relayed from the OFC once learning is crystallized^64,65^. Beyond the scope of the present study, more work is needed to better understand how value and latent task states signals in OFC and in visual regions dynamically interact during learning latent structures, and the division of labor between the regions in maintaining acquired latent structure over long time periods.

While in the current study context-dependent non-local updates promoted generalization performance in both humans and neural networks, these dynamics may also impair performance, for instance by over-generalizing or generalizing over irrelevant task features. For example, an overshoot in within-context pull effect in our task could potentially obscure behaviorally-relevant distinctions within a context. The precise influence on performance and decision-making is likely contingent on additional factors that govern non-local generalization over latent representations, as well as on how behavioral generalization is operationalized and measured experimentally. Investigating which conditions render non-local representation learning detrimental for decision-making is a potential novel avenue for explaining biases in decision-making. Would higher urgency during learning, for instance, induce larger co-updates, leading to overgeneralization?

Our study has several limitations, including moderate effect sizes, only correlational links to behavior, and lack of evidence for shifts driven by single feedback events. In our experimental design, irrelevant features remained fixed between the pre- and post-feedback estimation trials of each learning segment, which limited our ability to measure non-local updates structured by task-irrelevant dimensions. However, our network simulations suggest that the structure of non-local updates can be diagnostic of whether the true task structure is being learned. Future experiments varying task-irrelevant dimensions could test this directly.

The current findings are also useful when considering one prominent difference between modern AI systems and human brains: while parameter learning in artificial agents is restricted to a training phase and then fixed during deployment, learning in humans is an open-ended process ^66^. Understanding how the brain ceaselessly incorporates experiences to continuously update and reshape latent structures can provide insights into enabling artificial agents that are also open-ended, continual learners.

In sum, the findings demonstrate rapid, non-local dynamics of representation learning that reflect generalization over learned latent structures — a characteristic which can expedite learning and is essential for fast adaptive behaviour. Their presence in both the human OFC and deep networks offers a mechanistic starting point for understanding how abstract knowledge is continuously reshaped through, and in turn guides, decision making.

## Methods

This study was approved by the ethics committee of the German Psychological Society (DGP, reference: SchuckNicolas2022-07-05-WV) and complies with all relevant ethical regulations.

### Participants

A total of 64 right-handed, neurologically healthy participants with normal vision took part in this study. All participants gave written informed consent before participation. Participants were recruited using the participant database of the Max Planck Institute for Human Development. The first two participants who were scanned were excluded because they played an older version of the experiment, and one participant was excluded due to excessive motion. 10 additional participants were excluded since they did not reach performance criteria on learning (see below). A total of 51 participants (mean age 26 ± 4.5, 36 females) were therefore included in all analyses.

Performance criterion was a priori defined based on the accuracy on the last learning block (block 11). Participants having 5 errors or more in this block (<80% correct trials), were classified as unsuccessful learners and were excluded from all analyses. Of note, mean accuracy of unsuccessful learners on the last block was well below the threshold (63%), ranging from 46% to 75%.

### Experimental procedure

The experiment consisted of 3 scanning sessions across 2 subsequent days, two performed on day 1 and the third on day 2. The current study involved analyzing data from day 1, designed to test non-local representation learning. Two exceptions are that (i) visual localizers used to functionally define visual ROIs were collected on day 2 and (ii) the first estimation block collected on day 2 was included in the control analysis for non-local updates in the absence of local feedback (see section below). Upon arrival, participants completed on-screen task instructions, including practice trials (with occluded reward feedback), followed by comprehension check questions. They then completed two fMRI sessions (∼90 min each), separated by a ∼40 min break. Session 1 consisted of a pre-learning estimation block and 6 learning blocks. Session 2 consisted of 5 additional learning blocks and a final post-learning estimation block. The task and design are described in detail below. The task was implemented in Matlab R2020a using Psychtoolbox v.3. Imaging data was preprocessed using fMRIPrep. All analyses were carried out with Python.

### Task Design

The task was a partially-observable reinforcement learning task, framed as a cover story of training to be a realtor. Learning was achieved solely through learning trials, which comprised the bulk of the experiment. Each learning trial began with an image of a client (1 sec), followed by an ∼3 sec inter-stimulus interval (ISI) on average and two simultaneously presented house images (2.5 sec). The participant had to select, with a key press, which of the houses to offer to the previously observed client. Once a response was given, the central fixation changed its color from black to red for the remaining presentation time. A feedback displaying the number of points earned (0–100) appeared after a ∼3 sec ISI. If no response was recorded within 2.5 seconds, a prompt notifying the participants they were too slow in responding appeared instead of the reward feedback. A central fixation circle was displayed throughout the experiment (aside from during reward feedback), and participants were instructed to maintain their gaze centrally on the fixation. Inter-trial intervals and ISIs were always drawn from a truncated exponential distribution (mean = 3 sec, range 2–7 sec; mean value verified per block).

Participants’ goal was to maximize cumulative points, by learning the set of preferences different clients have towards different houses. They were told that not all client or house features would necessarily be relevant, and that their total points would be converted into a monetary bonus of up to 40 euros (maximal bonus possible for the first 2 sessions included in the current study was 26 euros).

Stimuli comprised 8 client images and 8 house images in total (Fig. 1c). Client images varied along 2 binary features: gender (woman/man) and eyewear (glasses/no glasses), with only one being task-relevant and yielding only two task-relevant client types (i.e., contexts). Each of the four resultant classes included two specific face identities, resulting in 8 images overall. House images varied along 3 binary features: house material (stone/wood), location (woods/mountains) and exterior facility (pool/swing), with only 2 being task-relevant and one out of the two being more important than the other (primary and secondary house features). The specific relevant features were assigned under one of two possible solutions, counterbalanced across participants: For the first solution, eyewear defined the client type, while location and house material were the primary and secondary task-relevant house features, respectively. For the second solution, gender defined the client type, while house material and exterior facility were the primary and secondary relevant house features, respectively. House stimuli were custom-designed for this study by a graphic designer. Face stimuli were synthetic, AI-generated faces obtained from Generated Photos (Generated Media, Inc., https://generated.photos) under a paid license.

State values were computed as a weighted linear sum of task-relevant house features (weights: ±3 for the primary feature, ±1 for the secondary feature), multiplied by a client-type weight (−1 or +2).

To sample non-local representational shifts, learning trials were interleaved with estimation trials. An estimation trial began with a client image (1 sec), followed by a ∼3 sec ISI and a single house display (1 sec). Participants were instructed to estimate the value of the client–house pair presented in a given estimation trial. To ensure they performed the task, in 40% of estimation trials a numerical probe appeared after another ∼3 sec ISI, and the participant had to indicate whether the value of the client-house pair was higher or lower than the number appearing on the screen (possible probes: 30/50/70). The probe appeared for 2 seconds regardless of response time, and turned red upon response collection. Importantly, no feedback was provided on these trials and participants could not learn from them.

The experiment included a sequence of 11 learning blocks, preceded and followed by a single pre- and post-learning estimation block. Learning blocks were structured as follows: Each learning block began and ended with a learning segment, with 16 learning trials in between. Learning segments included 4 pre estimation trials, 4 learning trials, and 4 post estimation trials. Altogether, each learning block included 24 learning trials, sampling every combination of task-relevant features once (2 possible client types X 4 possible left house types X 3 possible right house types). With regards to estimation trials, each learning block consisted of 16 estimation trials sampling each relevant client-house combination (i.e., state) twice, once in the pre phase of a learning segment and once in its post phase. Irrelevant features were kept fixed across the pre and post trials of each state. In the second learning segment of each block, we added an additional (fifth) estimation trial in the pre- and post segments which, by contrast to the other 4 states, was presented during the 4 interleaving learning trials, yet these estimation trials (22 in total per participant) were excluded from all non-local co-update analyses. The specific states assigned to the two learning segments per block rotated over 3 possible designs with a fixed order, counterbalancing 3 possible orders across participants. Regardless of the specific assignment per block, each learning block included one sample of non-local shifts for each of the 8 states.

Two separate cycling schemes, one for learning trials and one for estimation trials, were used to assign irrelevant features per trial. For learning trials, each of the 24 states had 8 possible irrelevant variants (3 irrelevant binary features). Irrelevant feature variants were assigned to states using a cyclic counterbalancing scheme, such that each state was paired with each of its 8 irrelevant feature variants exactly once every 8 blocks: variant assigned = (state id % 8 + shift) % 8, where shift increments by 1 after each learning block. The mapping of irrelevant feature combinations to variant indices was shuffled per participant, constrained so the irrelevant client feature alternates regularly across repetitions of the same state. For estimation trials, each of the 8 states had 4 irrelevant variants (2 irrelevant binary features). All estimation trials within the same learning segment used the same irrelevant variant, which incremented on each learning segment (i.e., twice per block, cycling through 4 options).

### Behavioral modeling

Linear approximation models consisted of a binary input vector ***x*** that encoded information about the inputs, and was connected to a single output node y via a weights vector W: *y* = *W**x***ᵀ. Weights were initialized randomly from a normal distribution *(mean= 0, SD = 0.25*). Because inputs were one-hot encoded, each value estimate initially equals a single weight. We therefore used the same initialization scale for both models, so that they started with identical uncertainty in their value estimates and differed only in their state representation. On every trial, the model received the two input vectors corresponding to the left and right options and predicted a value for each. The model then selected the same option as the participant. Upon receiving reward feedback on the selected option, weights were updated according to the Rescorla-Wagner rule:

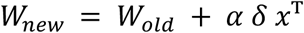

where ***δ*** is the prediction error: *δ* = *y* − *y*5, ***α*** is the learning rate, and ***x*** is the one-hot input vector of the chosen option.

To evaluate model fit, predicted values on each trial were converted to choice probabilities via a softmax function, and the negative log-likelihood (NLL) of the participant’s chosen option was averaged across all trials. Two free parameters, learning rate (***α***) and softmax inverse temperature (*β*), were estimated per participant via a grid search over 9 linearly spaced values per parameter (***α*** ∈ [0.1, 1]; *β* ∈ [1, 20]), selecting the parameter combination minimizing total NLL. Because both models had the same number of free parameters and differed only in their input representation, model fit was compared directly using NLL.

Input vectors in the perceptual model consisted of 64 nodes, each corresponding to a unique client-house combination. Input vectors in the compressed model consisted of 8 nodes, collapsing across the 8 variations in irrelevant client and house features per latent task state.

### MRI data acquisition

Visual stimuli were projected onto a screen via a computer monitor. Participants indicated their choice using an MRI-compatible button box using their right hand. Two horizontally-positioned buttons were used for left/right choices during learning trials. Two vertically-positioned buttons were used for more/less choices during estimation trials probes.

All MRI data were acquired using a 32-channel head coil on a research-dedicated 3-Tesla Siemens Magnetom TrioTim MRI scanner (Siemens, Erlangen, Germany) located at the Max Planck Institute for Human Development in Berlin, Germany. For the functional scans, whole-brain images were acquired using a segmented k-space and steady-state T2*-weighted multi-band (MB) echo-planar imaging (EPI) single-echo gradient sequence that is sensitive to the BOLD contrast (sequence specification: 80 slices in interleaved ascending order; anterior-to-posterior (A–P) phase-encoding direction; TR = 1750 ms; echo time (TE) =28.8 ms; voxel size = 1.75 × 1.75 × 1.75 mm; matrix = 96 × 96; field of view (FOV) = 196 × 196 mm; flip angle (FA) = 71°; distance factor = 0%; MB acceleration factor 4). Slices were tilted for each participant by 20° backwards relative to the Anterior Commissure - Posterior Commissure (AC-PC) axis, to improve the quality of fMRI signal from the orbitofrontal cortex^67^. Session 1 and 2 included 7 and 6 functional runs, respectively (learning and estimation blocks lasted for ∼9 and ∼10 minutes, respectively). For each functional run, the task began after the acquisition of the first four volumes to allow for scanner equilibrium. In each session, two short acquisitions with six volumes each were collected using the same sequence parameters as for the functional scans but with varying phase-encoding polarities, resulting in pairs of images with distortions going in opposite directions between the two acquisitions (also known as the *blip-up/blip-down* technique). From these pairs the displacement maps were estimated and used to correct for geometric distortions due to susceptibility-induced field inhomogeneities as implemented in the fMRIPrep preprocessing pipeline^68^. High-resolution T1-weighted (T1w) anatomical Magnetization Prepared Rapid Gradient Echo (MPRAGE) sequences were obtained from each participant in each session to allow registration and anatomical parcellation (sequence specification: 192 slices; TR = 1900 ms; TE = 2.52 ms; FA = 9°; inversion time (TI) = 900 ms; matrix size = 192 × 256; FOV = 192 × 256 mm; voxel size = 1 × 1 × 1 mm).

Respiration and pulse signals were recorded throughout each scanning session using a pneumatic respiration belt and pulse oximetry probe placed on the left index finger, respectively. Eye movements were recorded continuously at 1000 Hz via the EyeLink II system (SR Research).

### MRI data preprocessing

Functional and anatomical scans were preprocessed using *fMRIPrep*^68^ version 23.1.4, implementing the following routines:

#### Anatomical data

A total of 3 T1-weighted (T1w) images were found within the input BIDS dataset. All of them were corrected for intensity non-uniformity (INU) with N4BiasFieldCorrection^69^, distributed with ANTs^70^. The T1w-reference was then skull-stripped with a *Nipype* implementation of the antsBrainExtraction.sh workflow (from ANTs), using OASIS30ANTs as target template. Brain tissue segmentation of cerebrospinal fluid (CSF), white-matter (WM) and gray-matter (GM) was performed on the brain-extracted T1w using fast (FSL (version unknown),^71^). An anatomical T1w-reference map was computed after registration of 3 T1w images (after INU-correction) using mri_robust_template (FreeSurfer 7.3.2,^72^).

Brain surfaces were reconstructed using recon-all (FreeSurfer 7.3.2, RRID:SCR_001847, ^73^), and the brain mask estimated previously was refined with a custom variation of the method to reconcile ANTs-derived and FreeSurfer-derived segmentations of the cortical gray-matter of Mindboggle (RRID:SCR_002438,^74^). Volume-based spatial normalization to two standard spaces (MNI152NLin6Asym, MNI152NLin2009cAsym) was performed through nonlinear registration with antsRegistration (ANTs), using brain-extracted versions of both T1w reference and the T1w template. The following templates were selected for spatial normalization and accessed with *TemplateFlow* (23.0.0, ^75^): *FSL’s MNI ICBM 152 non-linear 6th Generation Asymmetric Average Brain Stereotaxic Registration Model* [^76^, RRID:SCR_002823; TemplateFlow ID: MNI152NLin6Asym], *ICBM 152 Nonlinear Asymmetrical template version 2009c* [^77^, RRID:SCR_008796; TemplateFlow ID: MNI152NLin2009cAsym].

#### Functional data

For each of the 13 BOLD runs per participant (across the 2 sessions), the following preprocessing was performed. First, a reference volume and its skull-stripped version were generated using a custom methodology of *fMRIPrep*. Head-motion parameters with respect to the BOLD reference (transformation matrices, and six corresponding rotation and translation parameters) are estimated before any spatiotemporal filtering using mcflirt (FSL,^78^). The estimated *fieldmap* was then aligned with rigid-registration to the target EPI (echo-planar imaging) reference run. The field coefficients were mapped on to the reference EPI using the transform.

BOLD runs were slice-time corrected to 0.834s (0.5 of slice acquisition range 0s-1.67s) using 3dTshift from AFNI (^79^, RRID:SCR_005927). The BOLD reference was then co-registered to the T1w reference using bbregister (FreeSurfer) which implements boundary-based registration^80^. Co-registration was configured with six degrees of freedom. Several confounding time-series were calculated based on the *preprocessed BOLD*: framewise displacement (FD), DVARS and three region-wise global signals. FD was computed using two formulations following Power (absolute sum of relative motions, ^81^) and Jenkinson (relative root mean square displacement between affines^78^). FD and DVARS are calculated for each functional run, both using their implementations in *Nipype* (following the definitions by ^81^). The three global signals are extracted within the CSF, the WM, and the whole-brain masks.

Additionally, a set of physiological regressors were extracted to allow for component-based noise correction (*CompCor*^82^). Principal components are estimated after high-pass filtering the *preprocessed BOLD* time-series (using a discrete cosine filter with 128s cut-off) for the two *CompCor* variants: temporal (tCompCor) and anatomical (aCompCor). tCompCor components are then calculated from the top 2% variable voxels within the brain mask. For aCompCor, three probabilistic masks (CSF, WM and combined CSF+WM) are generated in anatomical space. The implementation differs from that of Behzadi et al. in that instead of eroding the masks by 2 pixels on BOLD space, a mask of pixels that likely contain a volume fraction of GM is subtracted from the aCompCor masks. This mask is obtained by dilating a GM mask extracted from the FreeSurfer’s *aseg* segmentation, and it ensures components are not extracted from voxels containing a minimal fraction of GM. Finally, these masks are resampled into BOLD space and binarized by thresholding at 0.99 (as in the original implementation). Components are also calculated separately within the WM and CSF masks. For each CompCor decomposition, the *k* components with the largest singular values are retained, such that the retained components’ time series are sufficient to explain 50 percent of variance across the nuisance mask (CSF, WM, combined, or temporal). The remaining components are dropped from consideration. The head-motion estimates calculated in the correction step were also placed within the corresponding confounds file. The confound time series derived from head motion estimates and global signals were expanded with the inclusion of temporal derivatives and quadratic terms for each ^83^. Frames that exceeded a threshold of 0.5 mm FD or 1.5 standardized DVARS were annotated as motion outliers.

The BOLD time-series were resampled into standard space, generating a *preprocessed BOLD run in MNI152NLin6Asym space*. First, a reference volume and its skull-stripped version were generated using a custom methodology of *fMRIPrep*. The BOLD time-series were resampled onto the following surfaces (FreeSurfer reconstruction nomenclature): *fsnative*, *fsaverage*. All resamplings can be performed with *a single interpolation step* by composing all the pertinent transformations (i.e. head-motion transform matrices, susceptibility distortion correction when available, and co-registrations to anatomical and output spaces). Gridded (volumetric) resamplings were performed using antsApplyTransforms (ANTs), configured with Lanczos interpolation to minimize the smoothing effects of other kernels^84^. Non-gridded (surface) resamplings were performed using mri_vol2surf (FreeSurfer).

### fMRI single trial activity maps

fMRI analyses were carried on single trial beta maps modelled with the Least Squares Single (LSS) method, best suited to account for temporal overlaps given our fast event-related design. For learning and estimation trials, we analyzed the betas corresponding to the house pair / single house display, respectively. All analyses were performed in the participant’s native volumetric brain space.

All analyses were performed in each participant’s native volumetric brain space. fMRI analyses were carried on single-trial beta maps modelled with the Least Squares Single (LSS) method, best suited to account for temporal overlap given our fast event-related design ^85^. Each event within each learning/estimation trial (client, single house, house pair, reward and estimation probe) was modelled by a dedicated GLM, with one regressor modelling the event of interest and another regressor modelling all other events and trials. For learning and estimation trials, we analyzed the betas corresponding to the house pair and single house event in each trial, respectively. Each event-specific GLM was estimated with nilearn’s FirstLevelModel (SPM canonical double-gamma HRF, AR(1) noise prewhitening). Nuisance regression included 24 motion regressors (six rigid-body realignment parameters plus their temporal derivatives, squares, and squared derivatives), the mean white-matter and csf signal, and a 128 sec discrete cosine transform (DCT) high-pass filter, all extracted from the fMRIPrep confound output. Physiological noise was additionally regressed out using RETROICOR regressors ^86^ as implemented in TAPAS PhysIO toolbox ^87^ and derived from concurrently recorded pulse oximetry and respiratory signals (pulse Fourier order = 3, respiratory Fourier order = 4, pulse × respiratory interaction order = 1; 18 regressors per run), appended to the confound set of each run’s design matrix. Spatial smoothing was applied to the extracted single-trial beta maps and restricted to each ROI mask (5 mm FWHM Gaussian kernel; data were masked before smoothing so the kernel could not blend in signal from outside the ROI).

### ROIs definitions

OFC and hippocampus ROIs were defined anatomically per participant based on the FreeSurfer aparc+aseg segmentation carried out by fMRIPrep. The OFC comprised 3 bilateral regions from the Desikan-Killiany atlas ^42^: medial orbitofrontal cortex, lateral orbitofrontal cortex, and pars orbitalis (labels: 1014, 2014, 1012, 2012, 1019, 2019). The hippocampus comprised bilateral hippocampal labels from FreeSurfer’s subcortical segmentation (labels 17 and 53). Because the OFC and hippocampus are particularly prone to susceptibility-related signal dropout ^67^, we identified and excluded voxels suspected of signal dropout using an approach similar to SPM’s implicit masking. For each participant, we first computed a subject-level mean global intensity by averaging, across all runs, each run’s own mean BOLD signal within the brain mask. A voxel was then retained in a given run only if its own mean raw BOLD signal, averaged over time within that run, exceeded 30% of this subject-level reference. This criterion was applied independently in every run, and a voxel had to pass it in all runs to be included in the final ROI mask. Visual inspection of the voxels classified as dropout on participants’ brains confirmed that they were concentrated in edge areas typical for signal dropout in these regions. The mean dropout ratio across participants was 6.9% (SD = 2.0%, range: 2.3– 11.5%) in OFC and 2.3% (SD = 2.4%, range: 0–8.6%) in HPC.

FFA and VTC ROIs were defined functionally in individual participants based on a visual localizer scan collected on the second day of the study (during session 3). The visual localizer followed a standard block design and included seven visual categories: faces, houses, cars, instruments, corridors, patterns, and phase-scrambled images. Participants 501–513 performed an earlier version of the localizer consisting of eight, partially overlapping visual categories: objects, scrambled objects, faces, bodies, houses, and scenes. The scans were modelled using a general linear model implemented with nilearn’s FirstLevelModel class (5 mm FWHM spatial smoothing; AR(1) noise prewhitening), with a regressor per visual category. ROIs were then defined following typically used contrast maps within an anatomical mask comprising the fusiform, inferior temporal, lateral occipital, lingual, and parahippocampal gyri (Desikan-Killiany parcellation). FFA voxels were selected using the contrast faces > places, wherein places included the categories houses, scenes and corridors; VTC voxels were selected using the contrast all visual categories > scrambled objects (statistical threshold for both ROIs: t > 2.33, uncorrected; equivalent to p< .01, one-tailed).

### RSA-GLM analysis

Single trial betas for learning and estimation trials were independently z-scored per voxel and run to qualify a cross-run analysis. For modelling learning trials, pattern similarity between all cross-run single trial pairs in the early (blocks 1-6) and late (blocks 6-11) learning phase was estimated by Pearson’s r. Neural similarity was modelled using a linear mixed-effects model (LME):

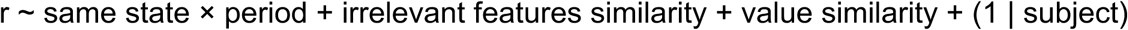

Same state regressor was binary (1: same, 0: different), and indexed trial pairs having the same two states comprising each trial, irrespective of the side in which houses were positioned or the irrelevant features. The irrelevant-feature regressor coded the proportion of irrelevant features shared by the two trials, out of the three irrelevant features present on each learning trial: the client feature, the left-house feature, and the right-house feature. The value similarity regressor was defined as 1 − |Δᵢ − Δⱼ| / Δmax, where Δᵢ and Δⱼ are the absolute reward differences between the two options on trials i and j, and Δmax is the maximum possible difference in the task, therefore it equals 1 for identical value differences and 0 for maximally different ones.

Estimation trials were modelled with the same approach, with two exceptions. First, these trials contained only two irrelevant features (an irrelevant client feature and a single irrelevant house feature as only one house was presented in estimation trials), so the shared proportion took values of 0, 0.5, or 1. Second, value similarity was based on the values of the single options in each trial, 1 − |vᵢ − vⱼ| / vmax, rather than on the difference between each trial’s value differences, as in learning trials.

To assess neural encoding independently per learning phase (denoted by single period significance stars in Fig. 3), each model was additionally re-fit separately for data in early and late periods, excluding the period term and its interaction with the same state regressor.

All models were fitted using the MixedLM function from the Python statsmodels library, optimised with L-BFGS (maximum 2000 iterations), with automatic fallback to Nelder-Mead (maximum 5000 iterations) upon non-convergence.

### Non-local representation learning analysis

A representational change vector was computed as the voxel-wise difference between the post- and pre-learning patterns elicited by the estimation trials pertaining to the same state (post minus pre). Co-update between two states was then defined as the Pearson correlation between their respective change vectors, reflecting whether their neural representations moved in similar/opposing directions across the intervening learning trials. For every participant, co-update values were computed for all pairwise state pairs in each learning segment, and were z-scored per participant and ROI. Co-update values were z-scored to center their distribution, results were unchanged when omitting this standardization step (see Fig. S3, ‘no z-score of co-update values’). Z-scored co-updates were then averaged across pairs within each learning segment separately for the two conditions of interest (Figure 4: same/different context; Fig. 5a: same/different house type among cross-context pairs) and subsequently averaged across learning segments to yield a single subject-level co-update estimate per condition and ROI. For cross-contexts value-dependent co-updates (Fig. 5b), no within learning segment averaging was needed, as each condition was assigned with a single co-update value (max/min value difference) per learning segment. Group-level statistics were derived from one-sample and paired-sample t-tests on subject-level means of the relevant co-update values.

### Control analysis for non-local updates in the absence of local feedback

To test whether non-local representational changes could arise without feedback, we repeated the co-update analysis on the two estimation-only blocks that followed learning blocks: the last block of day 1 and the first block of day 2. These blocks contained no learning trials, so non-local representational change between repeated presentations of a state cannot be driven by local feedback. Each block consisted of 64 trials in which every state was presented 8 times with different combinations of irrelevant features. Within each block, we built pseudo-learning segments that resembled the learning segments. Going through the block in order, the first presentations of 4 different states, two from each context, served as pre measurements. For each of these states, the post-measurement was its next presentation with the same irrelevant client and house features as at pre. As in the learning segments, irrelevant features were therefore held constant within each change vector. The next segment started on the trial after the last pre-measurement of the previous one, so that segments overlapped in time, but no trial contributed to more than one change vector This yielded a median of 11 pseudo-learning segments per participant. Intervals between pre- and post-measurements in the pseudo-learning segments were longer than in the learning segments (median 330 sec vs. 106 sec). All remaining steps were computed exactly as in the main non-local co-update analyses: voxel-wise z-scoring of patterns within each run, pairwise Pearson correlations between change vectors, and z-scoring of all co-update values per participant and ROI. Context- and value-dependent co-updates were computed within each segment, averaged across segments, and tested with paired t-tests (same vs. different context; maximal vs. minimal value difference), complemented by Bayes factors to quantify evidence in favor of a null effect (BF₀₁^88^). Bayes factors were computed with the python Pingouin package^89^, using the default Cauchy prior on effect size (scale = 0.707, i.e. 50% prior probability the effect lies between -0.707 and 0.707). Two participants had one missing estimation block, both had their remaining block included (N = 51). Excluding these two participants, who had the fewest segments (6 each; 9–13 for all others), did not change the results (all BF₀₁ > 5.6).

### Analytic derivation of non-local updates in deep linear networks

To characterize how feedback on one item non-locally shifts the representations of other items, we consider a linear network with two hidden layers and a single output node representing predicted value. To isolate shifts that are not attributable to input similarity, we assume that the inputs of different items are orthogonal.

The network maps an input x onto two hidden representations and a scalar output:

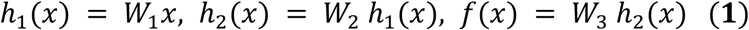

Feedback is given on a single item, the anchor (***x_a_***, ***y_a_***). The network minimizes the squared prediction error 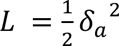, where ***δ***ₐ = ***y***ₐ − ***f***(***x***ₐ) is a scalar. A gradient descent step with learning rate ***η*** yields the following anchor-driven weights update:

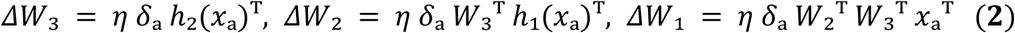

Each weight change is thus the outer product of the prediction error, ***δ***, propagated back to the layer’s output units, and the anchor’s activity at the layer’s input. Because the weights are shared across items, this update can also change the representations of items that received no feedback, which we refer to as probes. We define the non-local update of a probe ***x_i_*** at hidden layer ℓ as the resulting change in its representation, ***Δh*_1_**(***x_i_***).

### Non-local updates in the first hidden layer

The non-local update at the first hidden layer is obtained by applying *ΔW*_1_ (Eq. 2) to the probe’s input:

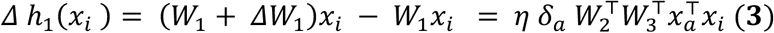

A probe’s first-hidden-layer representation therefore changes in proportion to its input similarity with the anchor. Under orthogonal inputs, 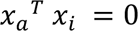, and feedback on the anchor produces no non-local updates at the first hidden layer, but Eq. 3 holds for arbitrary inputs and implies that first layer updates will always be fixed to input overlap regardless of learning.

#### Non-local co-updates in the first hidden layer

To characterize how feedback on the anchor shifts two probes relative to each other, we define their non-local co-update as the dot product of their non-local updates. From Eq. 3:

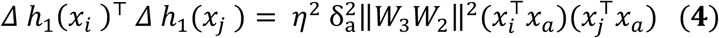

Because the term 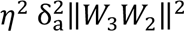 is a scalar independent of *i* and *j* that is the same for all non-local updates, this does not change the co-update similarity structure, which instead is related only to the input correlations between probes and targets. Hence the first hidden layer does not exhibit co-updates based on learned representational structure.

#### Non-local updates in the second hidden layer

The non-local update at the second hidden layer is *Δ* ℎ_2_(*x_i_*) = (*W*_2_ + *ΔW*_2_)(*W*_1_ + *ΔW*_1_) *x_i_* − *W*_2_*W*_1_*x_i_* . Because under orthogonal inputs *ΔW*_1_ *x_i_* = 0 (Eq. 3), this reduces to *ΔW*_2_ applied to the probe’s first-hidden-layer representation:

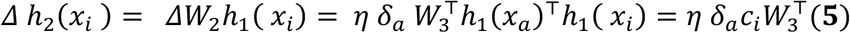

where *c_i_* = ℎ_1_(*x_i_*)^T^ℎ_1_(*x_a_*) is the similarity between the probe and anchor at the first hidden layer, which is acquired through learning, cᵢ is a probe-specific scalar, while δₐ is a probe-specific vector with dimensionality equal of that of the networks’ output, such the shared direction is W₃ᵀδₐ. Eq. 5 is exact and holds for any weights, i.e. at every point during training. Because the probe’s representation enters only through a scalar dot product, all probes in the second hidden layer are shifted along the same vector *W*_3_^T^, and differ only in the magnitude and sign of their shift.

#### Non-local *co-updates* in the second hidden layer

For the second hidden layer, the non-local co-updates are:

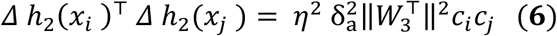

The co-update thus factorizes into two terms. The first, 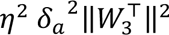, is the squared magnitude of the prediction error propagated back to the second hidden layer, which is shared by all non-local co-updates. It is positive and identical for all probe pairs, and therefore only scales the magnitude of the co-update. The second term, the product of the two probes’ learned similarities to the anchor *c_i_c*_j_ = ℎ_1_(*x_a_*)^T^ℎ_1_(*x_i_*)ℎ_1_(*x_a_*)^T^ℎ_1_(*x*_j_), is pair-specific and determines how the two probes shift relative to each other. Because all non-local updates lie along the same vector, each probe is shifted along it in the same direction as the anchor if its learned similarity to the anchor is positive, and in the opposite direction if it is negative. Consequently, two probes whose similarities to the anchor share the same sign, whether both positive or both negative, are shifted in the same direction (positive co-update), whereas two probes whose similarities have opposite signs are shifted in opposite directions, i.e. pushed apart (negative co-update).

Applying the same derivations deeper than the second hidden layer, directionality of non-local co-updates would gain degrees of freedom, beyond being pushed or pulled along a single direction.

### Neural networks simulations

#### Performance-matched neural networks

For the neural network setup, each trial was encoded as a two-step sequential input: at time step 1, a 10-dimensional vector encoding client features (4 first entries: 2 for relevant client feature, 2 for irrelevant client feature, rest zero-padded); at time step 2, a 10-dimensional vector encoding the features of one house option (6 last bits: 2 for primary house feature, 2 for secondary house feature, 2 for irrelevant house feature, rest zero-padded). The left and right options were encoded independently and passed through the same shared network to predict separate scalar value estimates.

The architecture consisted of a hidden recurrent layer (SimpleRNN, 32 units, tanh activation, no bias, termed here RNN layer), followed by a fully connected layer (16 units, ReLU, no bias, termed here FC layer), and a linear output layer (1 unit). Weights were initialized from truncated normal distributions (mean = 0, truncated at ±2 SD). For each layer’s weights, the standard deviation was a base value of √(2/(n_in + n_out)) multiplied by one of two gain parameters: the recurrent weights of the RNN layer were scaled by g_cdd_, whereas the input weights of the RNN layer and the weights of the FC and output layers were scaled by g_ef_. On each learning trial, the network estimated values for the left and right option, selected an action via softmax (temperature fixed at 1), received a scalar reward, and updated all weights via a single step of gradient descent (MSE loss, batch size = 1). The network updated its weights based on its own action choices rather than the participant’s, allowing it to develop an independent learning trajectory whose pace was matched to participants via parameter fitting (see below).

To obtain performance-matched simulations, we fitted three free parameters per participant: learning rate (η ∈ [0.01, 0.15]), the gain of the recurrent layer weights (g_cdd_ ∈ [0.01, 3]), and the gain of the remaining layers’ weights (g_ef_ ∈ [0.01, 3]). Fitting was performed via an exhaustive grid search with nine linearly spaced values per parameter, yielding 729 (9³) combinations. For each parameter combination, the network was trained with 10 random initializations. The best-fitting parameters were those minimizing the mean squared error between the network’s and the participant’s learning curves (trial-wise accuracy smoothed with a 15-trial moving average). A held-out 11th seed, withheld from the parameter selection process, was used to generate the final performance-matched network simulations for all subsequent analyses.

#### Non-local co-updates in neural networks

To quantify non-local representation learning in the performance-matched neural networks, we probed hidden-layer activations by presenting the same estimation trials inputs as in the experiment (i.e., immediately before and after interleaved learning segments) and logging their hidden activations. Change vectors were computed as the change in hidden-units activation between pre-and post-learning estimation trials of the same state (post − pre). Co-update was defined as the Pearson correlation between change vectors of two non-feedbacked states within the same mini-block, computed separately for same-context and different-context pairs, mirroring the fMRI analysis.

Prior to computing co-update values within a given learning segment, we applied two corrections to the pre- and post-learning representational geometries, both estimated from the same set of probed states, to remove global, non-specific representational changes. First, we mean-centered the pre- and post-learning representations separately by subtracting, from every state’s representation, the mean representation across the states probed in that segment, thus removing any global drift in representational space shared across all probed states^90^. Second, we applied an orthogonal Procrustes rotation^91^ (rotation and reflection only; no scaling), aligning the pre-learning geometry onto the post-learning geometry. This step removed any global rotation of the representational space that leaves pairwise geometry unchanged but can otherwise lead to spurious change-vector correlations^92,93^.

To assess whether the observed effects depended on the specific trial positions probed, we computed co-update traces under two sampling schemes, reported side by side throughout. In the empirical sampling scheme, hidden-layer activations were probed at the same trial positions as in the fMRI experiment and the co-update analysis was restricted to the four states probed at each learning segment. In the dense sampling scheme, activations were instead probed prior to, and immediately after, every learning trial, while excluding the two states presented in the learning trial from the co-update analysis.

For the empirical sampling scheme, performance on estimation trials was assessed using the same probe values and probe positions from the experiment itself. For the dense sampling scheme, every estimation trial included in the co-update analysis was assigned a pseudorandom (fixed seed) probe threshold (30/50/70).

#### Control analysis for impact of input overlap

Unlike the OFC, a context-dependent co-update effect was evident at the very first learning segment in both hidden layers of performance-matched networks (Fig. 7c,f). Based on our analytic solution of non-local updates in deep linear networks (see Methods section), we hypothesized this onset effect reflects greater visual similarity of within-versus cross-context input pairs, as the former always share the relevant client feature. To test this, and to ensure the prolonged effect is not driven by a greater visual overlap in same-context pairs, we applied the same co-update analysis on item pairs having the same versus different *irrelevant* context feature (i.e. irrelevant client feature). Given that the same asymmetry in visual overlap would apply when inputs are grouped by the identity of the irrelevant, rather than the relevant, context feature, any effect driven by visual overlap per se should emerge here as well. As shown in Fig. S6, an onset effect was evident for this grouping as well. However, it gradually decreased -rather than increased - with learning, demonstrating that the observed non-local context-dependent co-update effect cannot be accounted for by increased visual overlap for same context pairs.

#### Training with Oja’s rule

Oja’s rule was applied to the same two-layer recurrent network used for the performance-matched simulations (32-unit recurrent tanh layer, 16-unit fully connected ReLU layer, linear readout, and no biases), as well as the same input encoding, scalar reward and mean squared error objective as described above. The outer loop optimization for maximizing the fit to individual learning curves was achieved through a grid search, as described above for the gradient descent case, over learning rate, weights gain in the RNN hidden layer, weights gain in the FC hidden layer. The analysis was performed exactly as with the gradient descent networks, i.e. co-updates were computed from the change vectors of held-out states, after global drift subtraction and Procrustes alignment.

Applying the naive Hebbian update (Δwᵢⱼ = η · yᵢ · xⱼ) will typically result in divergent weights, so regularization needs to be added. This can be done by adding a term in the update rule that is designed to keep the weight norm small. Oja’s rule ^44^ is the canonical example of this, using a negative multiplicative weight term. We can write the vector of recurrent unit activations as y = tanh(Wx), where the vector x is the concatenation of the current input with the current hidden activation, the matrix W correspondingly contains the input weights in its first columns and the recurrent weights in its remaining columns, and the nonlinearity tanh (⋅) is applied elementwise. Oja’s weight update using the subspace rule ^94^ then takes the form:

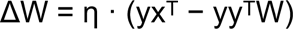

The second term ensures weights norms stay bounded. Because Oja’s rule is unsupervised (it does not depend on the target outputs y*), it is incapable of solving the task, and can only be used for learning a potentially better representation of the inputs. Thus, the linear readout was trained separately by gradient descent on the reward-prediction error. Since this layer has direct access to the outputs, using gradient descent here does not require backpropagation and is therefore local. All other layers were trained using Oja’s rule.

## Data availability

Anonymized behavioral and neuroimaging data will be made publicly available upon publication.

## Code availability

All code for neuroimaging analysis and simulations will be made publicly available on GitHub upon publication.

## Acknowledgments

N.W.S. was funded by the Federal Ministry of Education and Research (BMBF) and the Free and Hanseatic City of Hamburg under the Excellence Strategy of the Federal Government and the Länder, as well as a Starting Grant from the European Union (ERC-StG-REPLAY-852669). A.M.S. was funded by a Sir Henry Dale Fellowship from the Wellcome Trust and Royal Society (216386/Z/19/Z), the Sainsbury Wellcome Centre Core Grant from Wellcome (219627/Z/19/Z) and the Gatsby Charitable Foundation (GAT3755), and is a Canadian Institute for Advanced Research (CIFAR) Fellow. S.G. was funded by a Minerva Fellowship from the Minerva Stiftung and by the Zuckerman-CHE Postdoctoral Program.

## Supplementary figures and tables

**Table S1.**
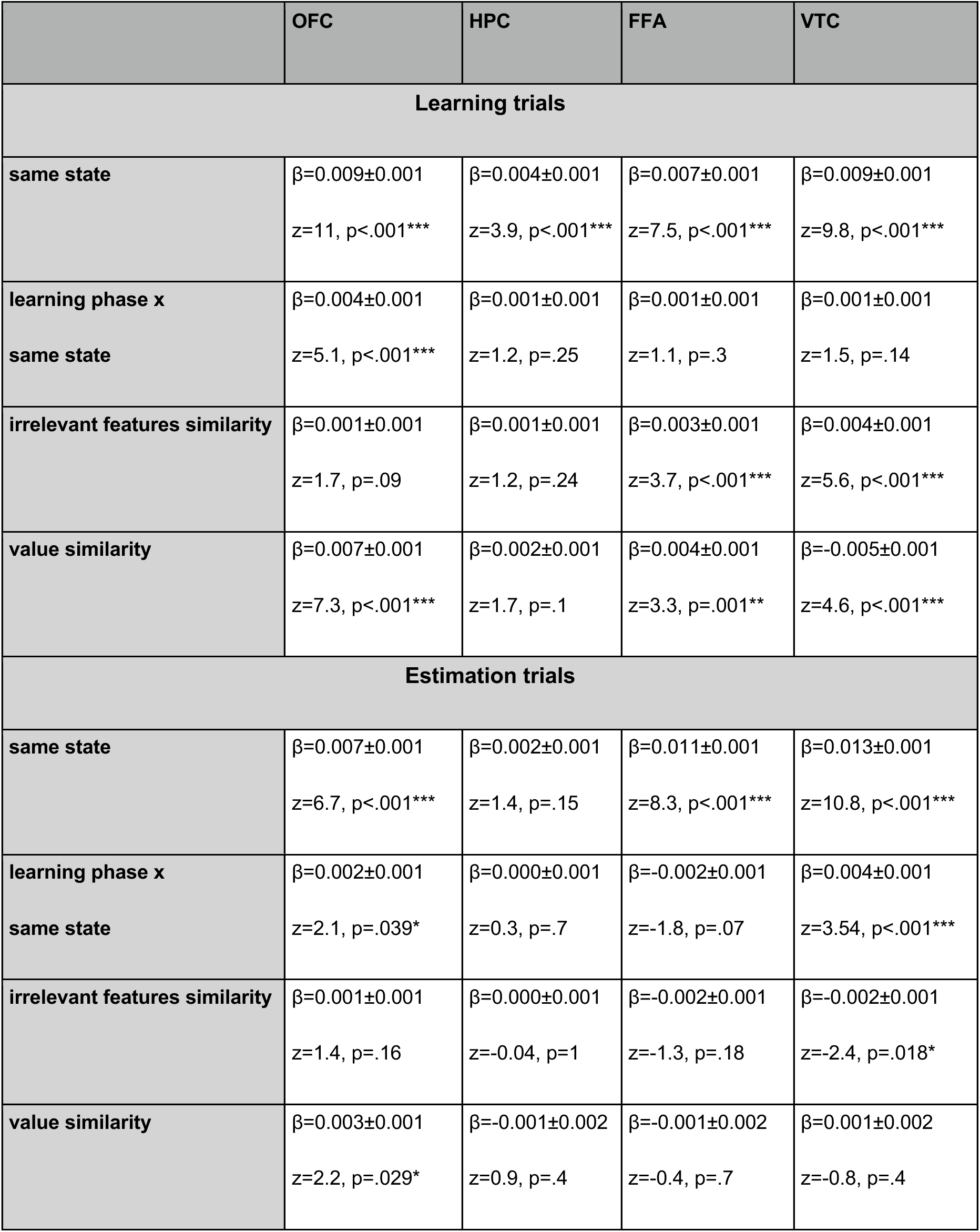
Summary of results obtained for the linear mixed-effects model depicted in Figure 3.

|  | **OFC** | **HPC** | **FFA** | **VTC** |
|---|---|---|---|---|
| **Learning trials** |  |  |  |  |
| **same state** | β=0.009±0.001<br>z=11, p<.001*** | β=0.004±0.001<br>z=3.9, p<.001*** | β=0.007±0.001<br>z=7.5, p<.001*** | β=0.009±0.001<br>z=9.8, p<.001*** |
| **learning phase x**<br>**same state** | β=0.004±0.001<br>z=5.1, p<.001*** | β=0.001±0.001<br>z=1.2, p=.25 | β=0.001±0.001<br>z=1.1, p=.3 | β=0.001±0.001<br>z=1.5, p=.14 |
| **irrelevant features similarity** | β=0.001±0.001<br>z=1.7, p=.09 | β=0.001±0.001<br>z=1.2, p=.24 | β=0.003±0.001<br>z=3.7, p<.001*** | β=0.004±0.001<br>z=5.6, p<.001*** |
| **value similarity** | β=0.007±0.001<br>z=7.3, p<.001*** | β=0.002±0.001<br>z=1.7, p=.1 | β=0.004±0.001<br>z=3.3, p=.001** | β=-0.005±0.001<br>z=4.6, p<.001*** |
| **Estimation trials** |  |  |  |  |
| **same state** | β=0.007±0.001<br>z=6.7, p<.001*** | β=0.002±0.001<br>z=1.4, p=.15 | β=0.011±0.001<br>z=8.3, p<.001*** | β=0.013±0.001<br>z=10.8, p<.001*** |
| **learning phase x**<br>**same state** | β=0.002±0.001<br>z=2.1, p=.039* | β=0.000±0.001<br>z=0.3, p=.7 | β=-0.002±0.001<br>z=-1.8, p=.07 | β=0.004±0.001<br>z=3.54, p<.001*** |
| **irrelevant features similarity** | β=0.001±0.001<br>z=1.4, p=.16 | β=0.000±0.001<br>z=-0.04, p=1 | β=-0.002±0.001<br>z=-1.3, p=.18 | β=-0.002±0.001<br>z=-2.4, p=.018* |
| **value similarity** | β=0.003±0.001<br>z=2.2, p=.029* | β=-0.001±0.002<br>z=0.9, p=.4 | β=-0.001±0.002<br>z=-0.4, p=.7 | β=0.001±0.002<br>z=-0.8, p=.4 |

**Figure S1.**
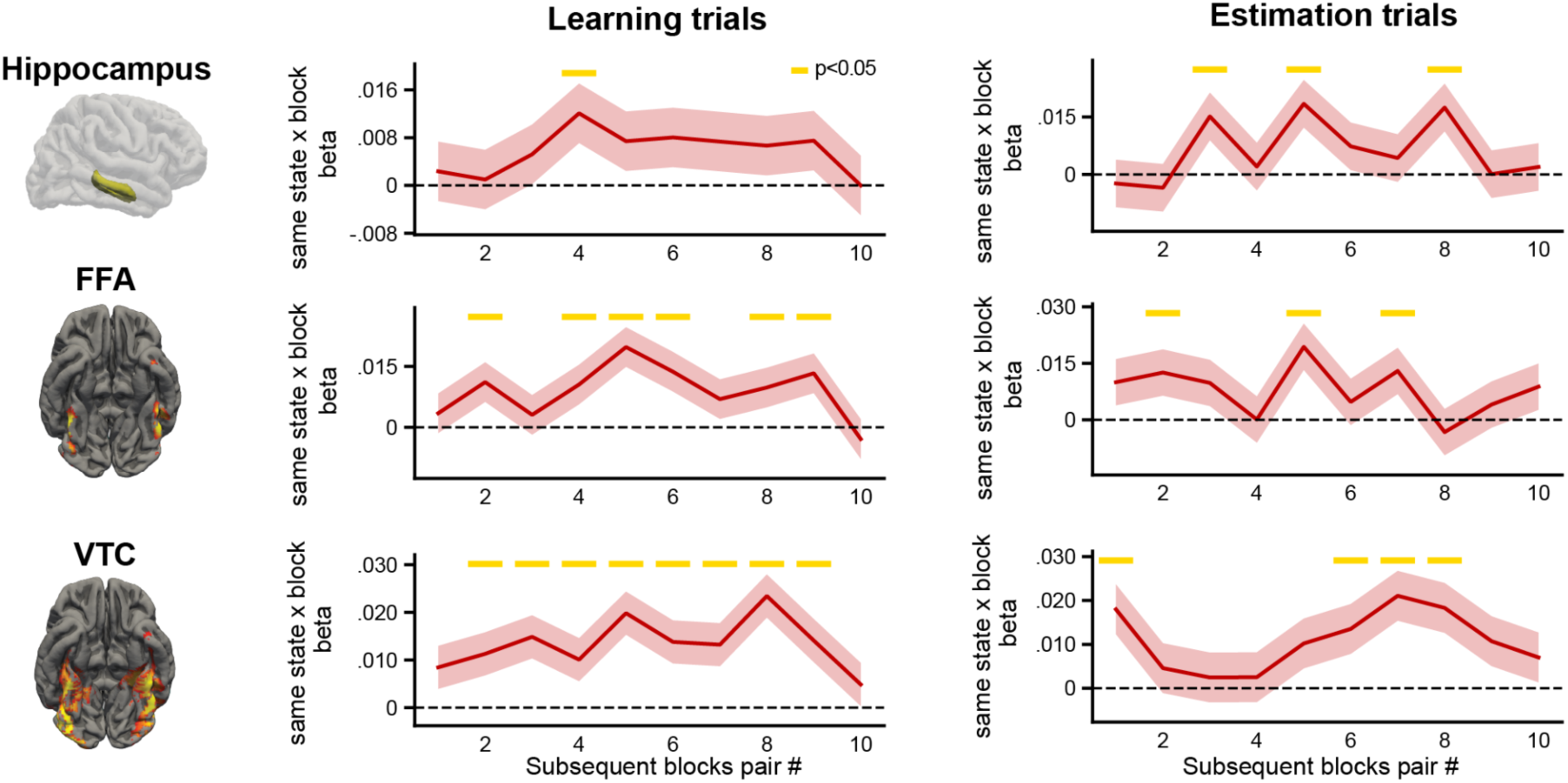
time-resolved state encoding over the course of learning in Hippocampus, FFA and VTC. Same analysis as presented in Figure 3a (bottom panel), applied to the three remaining ROIs.

**Figure S2.**
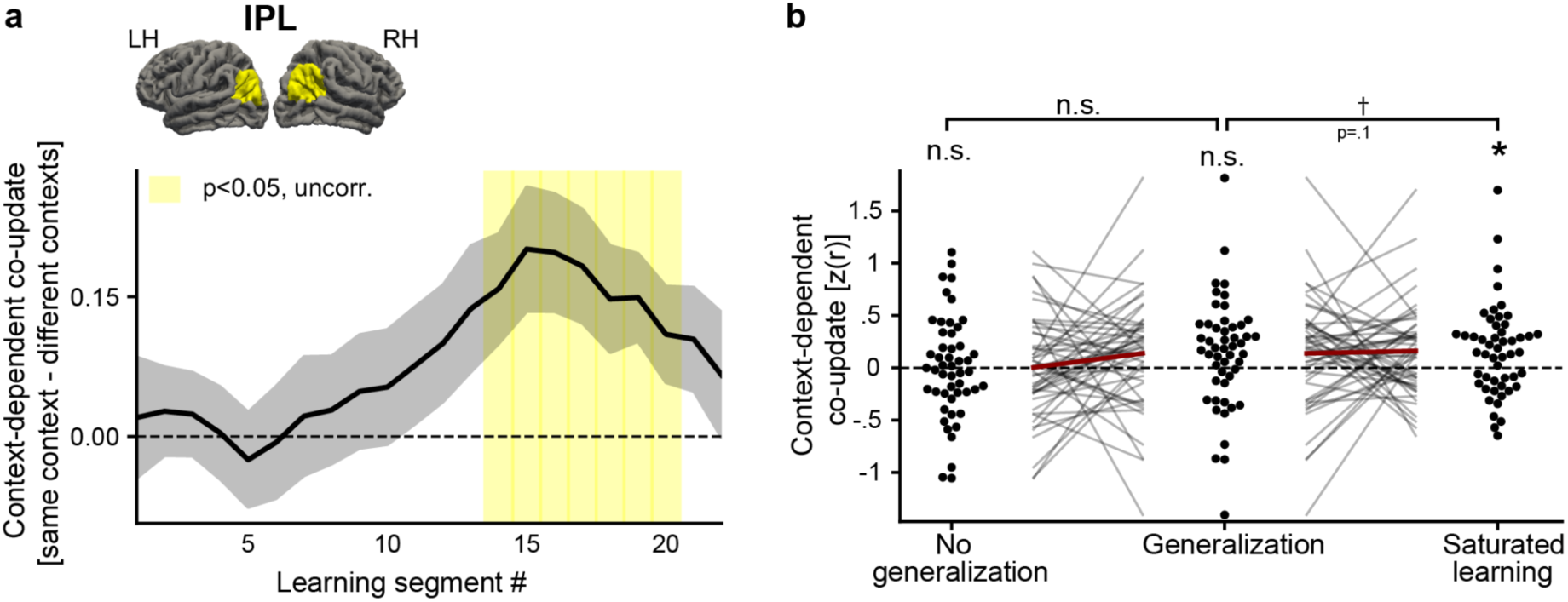
An exploratory analysis of context-dependent non-local co-updates and their link to performance in the Inferior Parietal Lobule (IPL). In an exploratory whole brain analysis of the mean context-dependent co-update effect, the IPL was the only region which showed an effect aside from the a-priori defined OFC (Fig. 4d). **a.** The time series of the effect for the IPL (equivalent to OFC’s analysis, Fig. 4c). **b.** Same analysis as in Fig. 6a, classifying the magnitude of non-local context-dependent co-updates by their concurrent performance changes, applied to IPL data. By contrast to the OFC, the IPL appears to show a later effect which does not coincide with representation learning, as also evident by a suggestive link to behaviour evident only during saturated learning.

**Figure S3.**
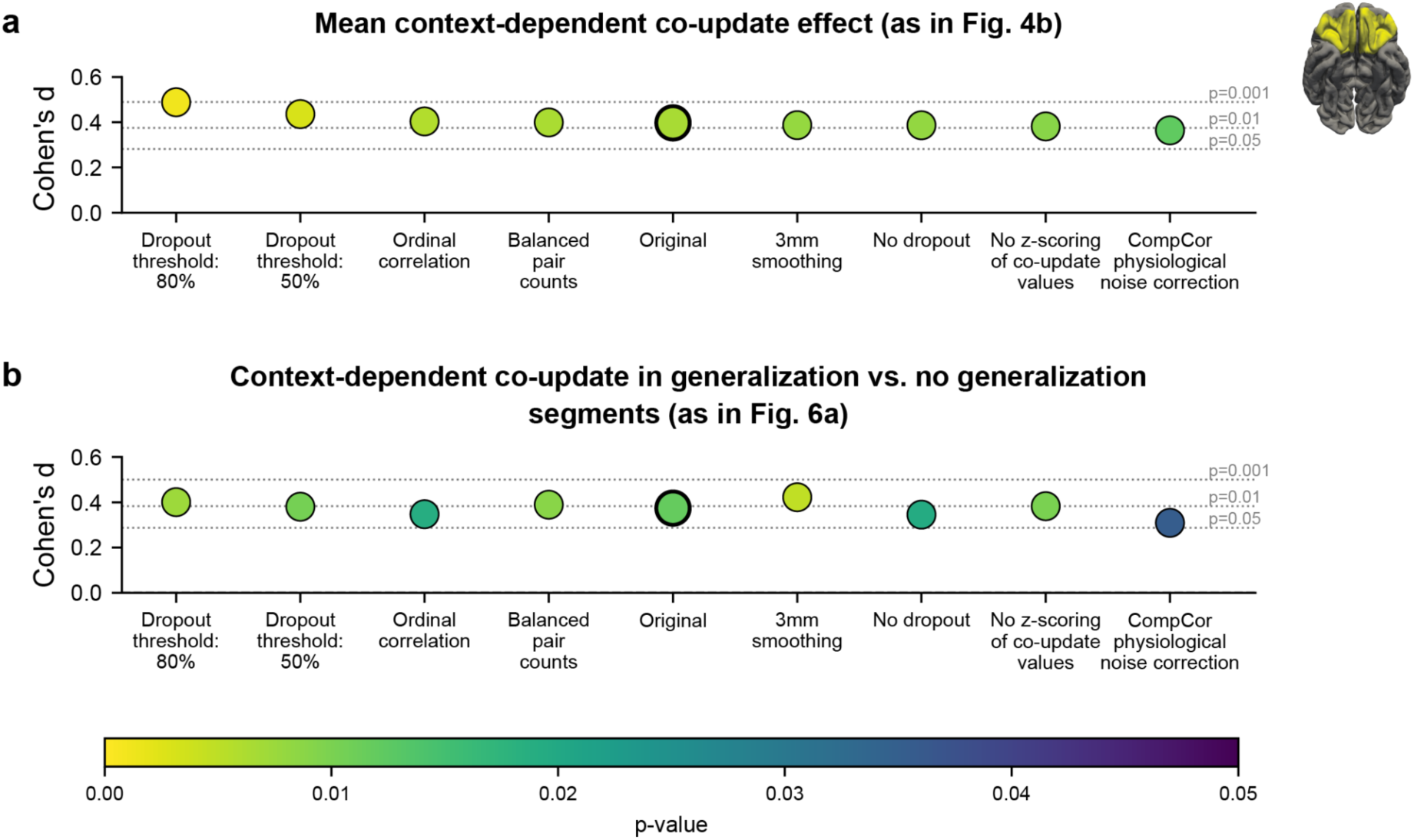
Robustness analysis of context-dependent co-updates in OFC and their link to generalization. **a.** Effect size (Cohen’s d) and p-value of mean context-dependent co-update in OFC across processing-pipeline variants [one-sample t-test against zero]. Variants are sorted in descending order by effect size; *Original* denotes the pipeline used in the main text. *Dropout threshold 80/50/0%*: threshold defining dropout voxels in OFC, following a similar procedure to the implicit brain masking implemented in SPM. The mean signal intensity across all TRs is computed for every voxel in the brain mask, and voxels whose mean fall below a given percentage of the global mean across all voxels are excluded (SPM’s default threshold is 80%). Higher thresholds are more stringent (higher dropout rates); 0% excludes no voxels. The original pipeline used 30%. *Rank-based correlation*: co-updates of change vectors estimated with Spearman’s ρ rather than Pearson’s r. *Balanced pair counts*: because each learning segment contained more cross-context than same-context co-update values by design, we subsampled cross-context values without replacement to match the same-context count (10 subsamples per segment) and ran all downstream steps on the mean co-update across matched subsamples. *No z-scoring*: omitting per-participant z-scoring of single co-update values. *CompCor physiological noise correction*: physiological noise modelled with PCA-based nuisance regressors from fMRIPrep (aCompCor, first 5 components from each of the white matter and CSF masks) instead of the original pipeline’s direct physiological recordings (respiration and pulse). **b.** The same variants applied to the effect linking generalization periods to increased context-dependent co-update relative to non-generalization periods (Fig. 6a, paired t-test). Variant order is fixed to that of panel a.

**Figure S4.**
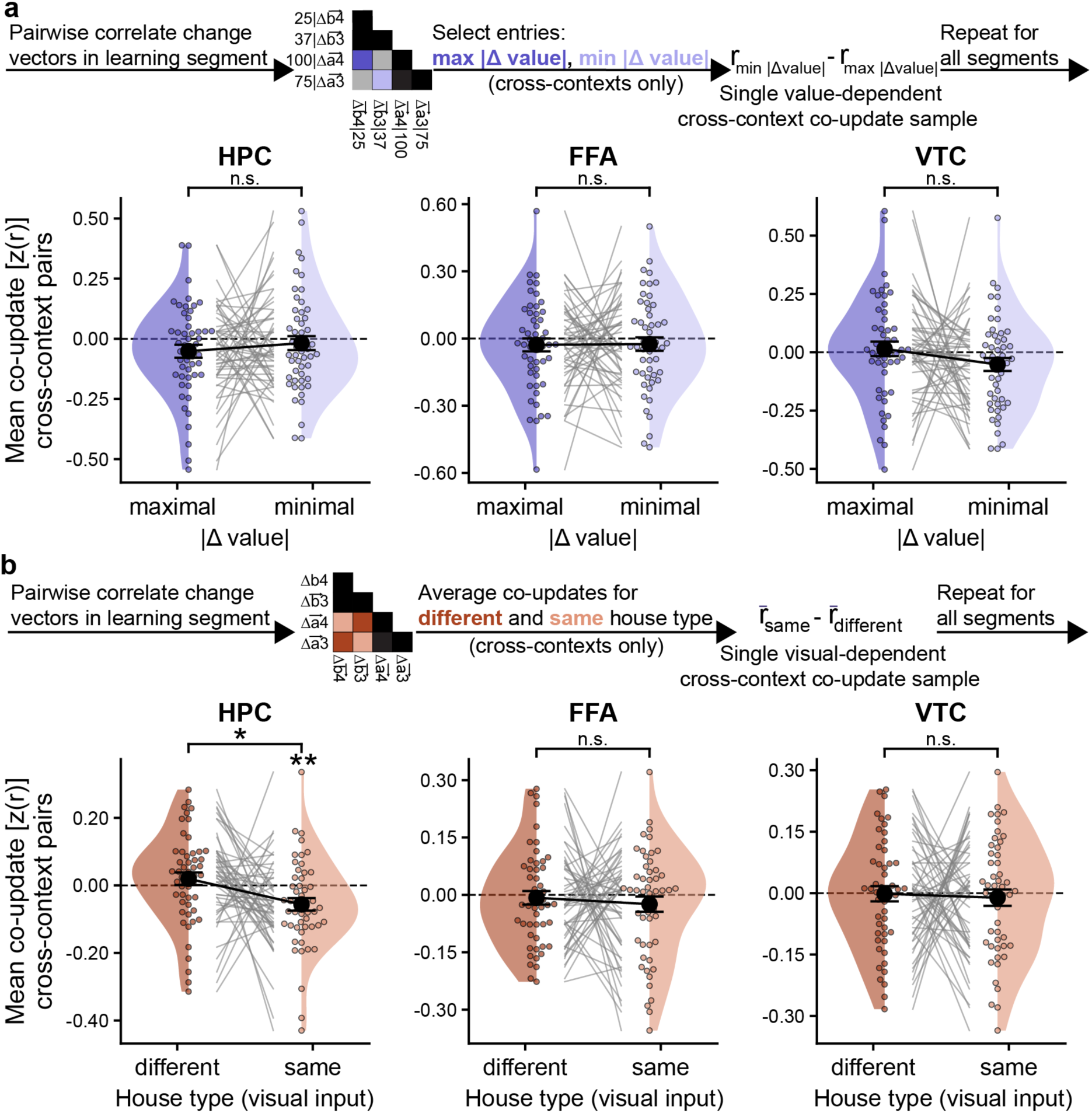
Visual- and value-dependent co-updates in remaining ROIs. Same analysis as in Fig. 5, applied to the remainder of the initially inspected ROIs that did not show a dynamical representation learning effect. Hippocampus showed a significant effect demonstrating greater representational divergence for cross-context state pairs with identical visual inputs.

**Figure S5.**
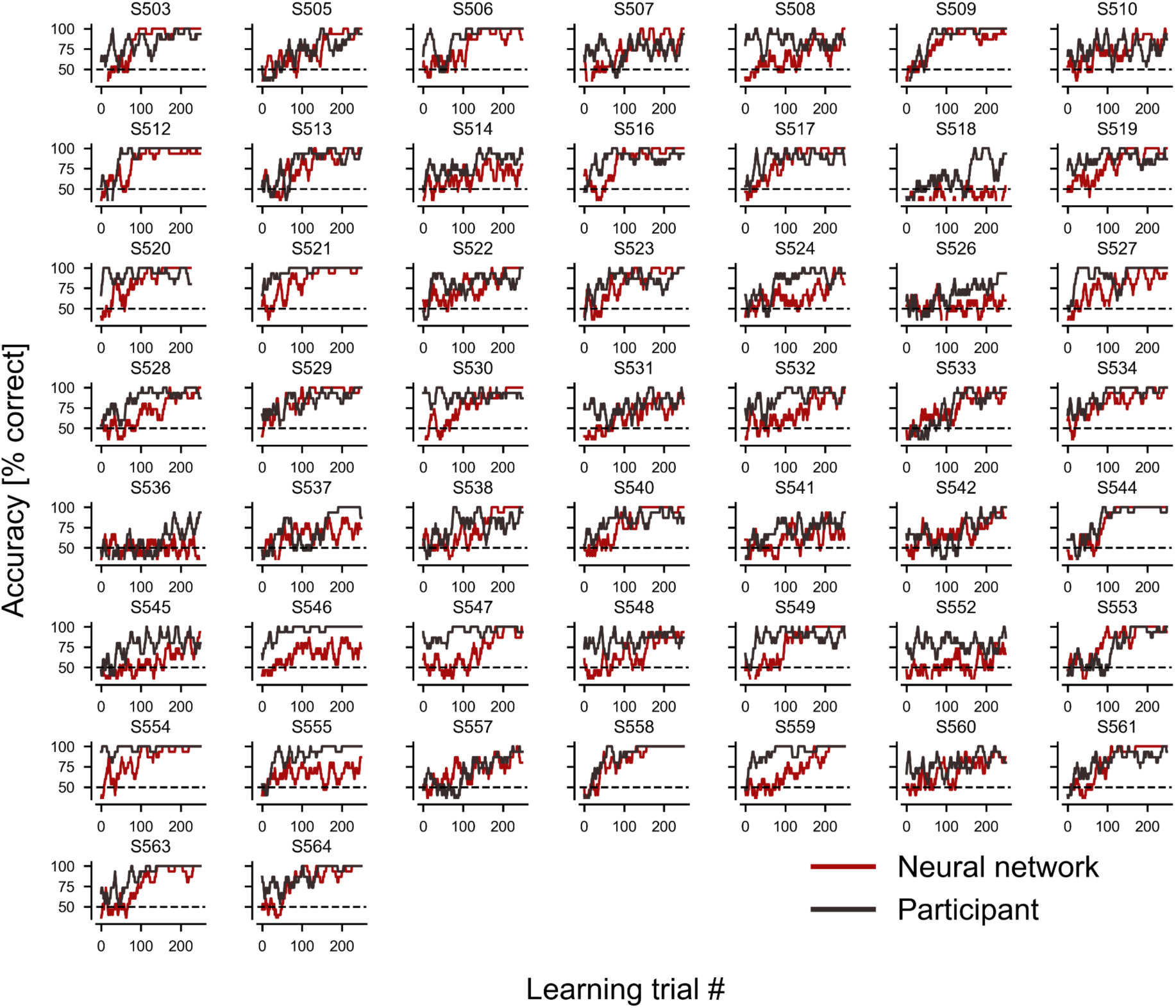
Learning curves of individual participants and fitted neural networks. Black: learning curves of individual participants depicting accuracy in learning trials over the course of 11 learning blocks (264 trials). Dark red: learning curve of neural networks trained on the exact same curricula as participants and optimized to best fit participants’ learning curves. Optimization performed over 10 random seeds and final simulation data obtained from an independent, held out seed). All curves are smoothed with a running average (window size = 15 learning trials).

**Figure S6.**
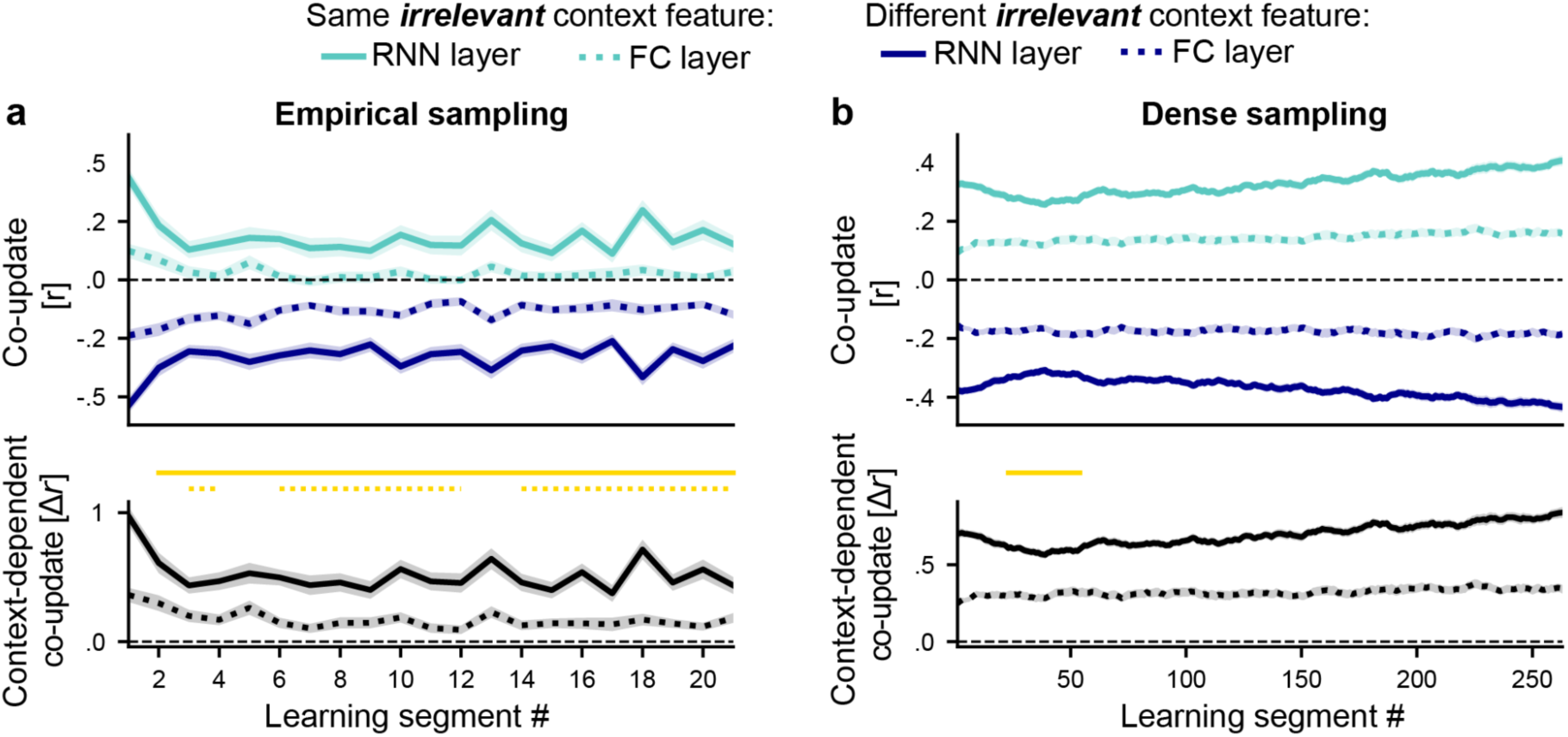
Impact irrelevant input overlap on non-local representation learning in neural networks. Same analysis as in Fig. 7c,f, except that co-updates were grouped by whether items shared the *irrelevant* rather than the relevant context feature. As with the relevant context feature, this grouping produced greater visual overlap for items sharing the feature, also evident in the onset difference between co-updates for same versus different irrelevant visual feature. In contrast to the relevant context feature, however, this difference was attenuated rather than amplified over the course of learning. This rules out visual overlap as an explanation for the gradual increase in non-local context-dependent co-updates observed in hidden layers of performance-matched networks. Yellow lines denote learning segments showing a significant decrease in irrelevant context-dependent co-update relative to the first learning segment (paired t-tests across networks, FDR-corrected at α=.05 within each layer; solid = RNN layer, dotted = FC layer).

**Figure S7.**
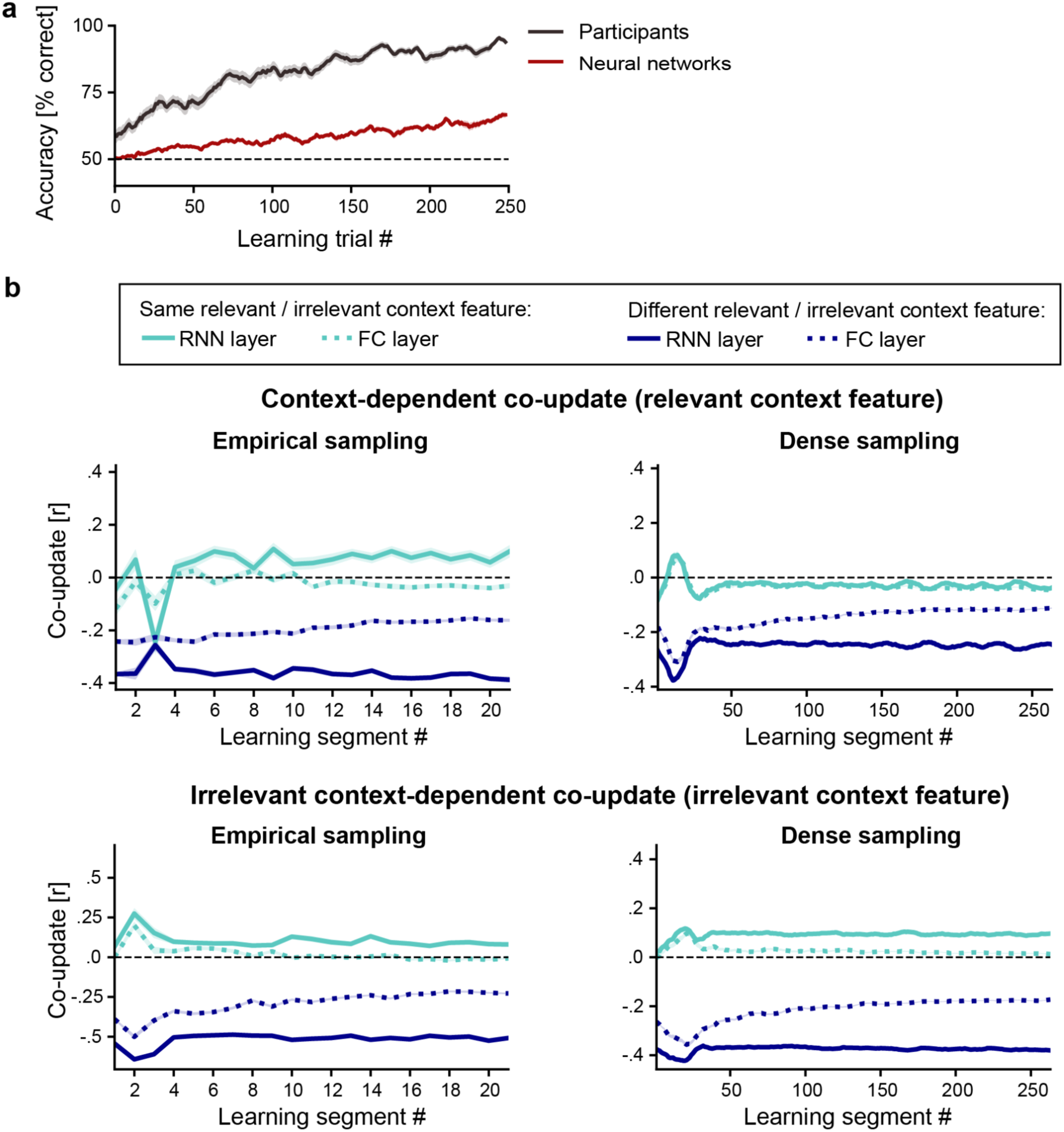
Accuracy and non-local co-updates in networks trained with Oja’s rule. **a.** Accuracy over learning segments for networks whose weights were updated with the Hebbian-like Oja’s rule, with hyperparameters optimized to fit participants’ learning curves. Networks did not reach human-level performance: because the Oja’s rule learns only from input correlations and incorporates no prediction-error signal, it cannot learn which features are relevant to the task. **b.** Non-local co-updates over learning segments for the task-relevant (top) and task-irrelevant (bottom) context feature. In both cases, co-updates for item pairs with greater input overlap on average emerged early and remained constant throughout learning, irrespective of task relevance. Thus, error-driven learning was required both to solve the task and to produce the relevance-specific, gradually increasing co-updates observed in the main networks.

